# Sirtuin 1 Activation Mitigates Murine Vasculitis Severity by Promoting Autophagy and Mitophagy

**DOI:** 10.1101/2025.09.04.671113

**Authors:** Asli E. Atici, Prasant K. Jena, Thacyana T. Carvalho, Benjamin L. Ross, Emily A. Aubuchon, Malcolm E. Lane, Angela C. Gomez, Youngho Lee, Shuang Chen, Timothy R. Crother, Moshe Arditi, Magali Noval Rivas

## Abstract

**BACKGROUND:** Sirtuin 1 (SIRT1), a NAD^+^-dependent protein deacetylase, regulates cardiovascular inflammation by modulating cellular stress, inhibiting NLRP3 activation, and promoting the clearance of damaged mitochondria. However, its precise role in the pathogenesis of Kawasaki disease (KD), a pediatric systemic vasculitis and the leading cause of acquired heart disease in children, remains unclear.

**METHODS:** Using the *Lactobacillus casei* cell wall extract (LCWE) murine model of KD, we evaluated the severity of vasculitis in mice supplemented with NAD^+^ precursors, as well as transgenic mice overexpressing SIRT1, and mice with specific deletion of Sirt1 in vascular smooth muscle cells (VSMCs) and myeloid cells. Proteomics analysis was performed on the abdominal aortas of WT and SIRT1-overexpressing mice. We performed immunofluorescent staining of cardiovascular tissues to assess the expression of proteins related to the autophagy/mitophagy pathway and the pathogenic switch of VSMCs. Western blot analysis was performed on primary VSMCs and cardiovascular tissues to determine the impact of SIRT1 on autophagic flux. The production of pro-inflammatory cytokines was measured in bone marrow-derived macrophages and peritoneal lavage of transgenic mice using ELISAs.

**RESULTS:** SIRT1 expression was downregulated in cardiovascular lesions of LCWE-injected mice, which was associated with a significant reduction of circulating levels of nicotinamide. Supplementation of mice with NAD^+^ precursors or genetic overexpression of SIRT1 significantly reduced the development of LCWE-induced KD, while the specific deletion of *Sirt1* in VSMCs or myeloid cells exacerbated vasculitis. Proteomics analysis indicated impaired mitophagy/autophagy and the pathogenic synthetic switch of VSMCs in LCWE-injected mice, which was rescued with SIRT1 overexpression and associated with reduced production of proinflammatory cytokines.

**CONCLUSIONS:** This study reveals the presence of an impaired NAD^+^-SIRT1 axis in the pathogenesis of LCWE-induced KD vasculitis and the therapeutic potential of targeting this axis to reduce cardiovascular lesions and inflammation.

## INTRODUCTION

Sirtuins are a highly conserved family of nicotinamide adenine dinucleotide (NAD^+^)-dependent deacetylases with broad physiological functionality, such as transcriptional regulation, energy metabolism, inflammation, and stress response ^1^. The best characterized member of the family, Sirtuin 1 (SIRT1), deacetylates histones and non-histone proteins, condensing chromatin and silencing gene transcription ^2,3^. Through these functions, SIRT1 exerts a beneficial role across a broad spectrum of pathologies, from inflammatory to metabolic and cardiovascular diseases ^2,3^. Indeed, SIRT1 mitigates vascular remodeling, atherosclerosis, arterial stiffness, and aortic aneurysms ^2–4^. Hence, there is an emerging interest in SIRT1 as a therapeutic target for cardiovascular diseases.

Kawasaki disease (KD) is an acute febrile systemic vasculitis of unknown etiology, affecting infants and young children, and the leading cause of acquired heart disease worldwide ^5,6^. Around 25% of untreated children develop coronary arteritis and coronary artery aneurysms (CAAs), a rate that is reduced to 3-5% following high-dose intravenous immunoglobulin (IVIG) treatment ^5^. However, up to 20% of patients are IVIG-resistant and are at higher risk of developing CAAs, requiring additional therapies ^5^. Infiltrations of monocytes, macrophages, neutrophils, CD8^+^ T cells, and IgA^+^ plasma cells into the coronary artery wall are observed during KD ^7–10^ and occur concurrently with increased levels of circulating pro-inflammatory cytokines, such as tumor necrosis factor α (TNF-α) and interleukin (IL)-1β ^11–13^. Human and murine studies have highlighted the crucial role of IL-1 in the development of KD cardiovascular lesions, but an improved understanding of KD pathophysiology is necessary to find novel therapeutic targets ^14,15^. The *Lactobacillus casei* cell wall extract (LCWE)-induced murine model of KD vasculitis reproduces key pathological manifestations of human KD, including myocarditis, coronary arteritis, aortitis, and luminal myofibroblast proliferation, which result in coronary artery abnormalities, such as dilations and aneurysms ^16,17^. In addition to CAAs, systemic artery aneurysms are reported in human KD ^18–20^, and LCWE-injected mice also exhibit dilations in peripheral arteries, including the infra-renal part of the abdominal aorta ^21,22^. Activation of the NLR family, pyrin domain-containing 3 (NLRP3) inflammasome, IL-1β production, and infiltration of macrophages to the inflamed heart and abdominal aorta are hallmarks of LCWE-induced KD vasculitis progression ^22–24^. The genetic or antibody-mediated blockade of the IL-1 pathway significantly reduces the severity of LCWE-induced vasculitis and the development of abdominal aorta dilations, indicating a crucial role for the NLRP-3/IL-1 pathway in LCWE-induced KD pathogenesis ^16,22,23^. Therefore, the LCWE-induced KD model is an invaluable tool to gain mechanistic insights into KD pathogenesis ^17^.

Mitochondrial damage and impaired autophagy/mitophagy can activate the NLRP3 inflammasome by impairing oxidative phosphorylation (OXPHOS) and triggering the release of mitochondrial DNA (mtDNA) from the mitochondria, which acts as a danger signal ^25–27^. The dysfunctional clearance of damaged mitochondria, or mitophagy, has been observed in LCWE-injected mice developing vasculitis, and was associated with a decreased autophagic flux and the inhibition of autophagy by mammalian target of rapamycin (mTOR) activity ^28^. This subsequently leads to NLRP3 inflammasome activation and ROS accumulation ^28^. SIRT1 has been shown to inhibit inflammatory responses and to decrease NLRP3 inflammasome activation and IL-1β production ^29,30^, suggesting that its activation may counteract KD progression. Furthermore, SIRT1 decreases VSMC proliferation ^31,32^, which contributes to luminal myofibroblast proliferation (LMP) and coronary stenosis during KD ^33^. SIRT1 also exerts an anti-inflammatory effect by promoting autophagy and counteracting mitochondrial dysfunction and the accumulation of damaged mitochondria ^34–36^.

While SIRT1 promotes autophagy/mitophagy and has anti-inflammatory properties, its therapeutic potential for KD vasculitis remains unexplored. Here, we used the LCWE murine model of KD to investigate the impact of SIRT1 on the development of LCWE-induced KD vasculitis. We report that activating SIRT1 promotes autophagy/mitophagy, reduces ROS accumulation, decreases the production of bioactive IL-1β by macrophages, and prevents the pathogenic switch of VSMCs in LCWE-induced cardiovascular lesions. Overall, our results emphasize the critical role of SIRT1 in regulating LCWE-induced cardiovascular inflammation, highlighting the potential of targeting its activity as a novel therapeutic approach for KD.

## RESULTS

### NAD^+^ pathway metabolites and SIRT1 levels are decreased in LCWE-induced KD

As previously reported, a single injection of LCWE triggers the development of cardiovascular inflammation in wild-type (WT) mice (**Figure 1A-C**) ^16,21–24,28^. This inflammation is characterized by the infiltration of immune cells around the aorta (aortitis) and the coronary artery area (**Figure 1A**), as well as the development of dilations in the infrarenal part of the abdominal aorta (AA) (**Figure 1B, C**) ^16,21–24,28^. LCWE-induced vascular inflammation is an immune-mediated vasculitis, driven by IL-1β and involving the infiltration of innate and adaptive immune cells into cardiovascular lesions, resulting in vascular tissue damage ^16,21,22^. Since intense inflammatory processes impact host metabolism ^37^, we performed a metabolomic analysis of serum samples collected from WT mice at baseline (Day 0) and at 2 weeks post-LCWE injection (Day 14), to identify metabolic alterations linked to LCWE-induced KD vasculitis (**Figure S1A**). Principal components analysis (PCA) indicated distinct clustering patterns between the two groups (**Figure 1D**), and we identified 108 metabolites that were significantly different (*p*-value ≤ 0.05) between the two groups, with 82 being upregulated and 26 being downregulated 14 days after LCWE-injection when compared with the baseline timepoint (**Figure 1E**). LCWE injection and KD vasculitis development were associated with significant alterations in 9 pathways. Among these was the nicotinate and nicotinamide pathway, which was decreased in LCWE-injected mice compared with baseline (**Figure 1F**). We observed decreased levels of circulating nicotinamide (NAM) and nicotinamide N-oxide, precursors to NAD^+^, suggesting a potential deficit in NAD^+^ production by the salvage pathway (**Figure 1G** and **Figure S1B**). In addition, normalized scaled intensities of NAM, 1-methylnicotinamide, and nicotinamide N-oxide were significantly reduced after LCWE-injection compared with baseline (**Figure 1H**). The significant decrease in circulating NAD^+^ levels in LCWE-injected mice was confirmed in a separate cohort of PBS and LCWE-injected mice (**Figure 1I**).

**Figure 1.**
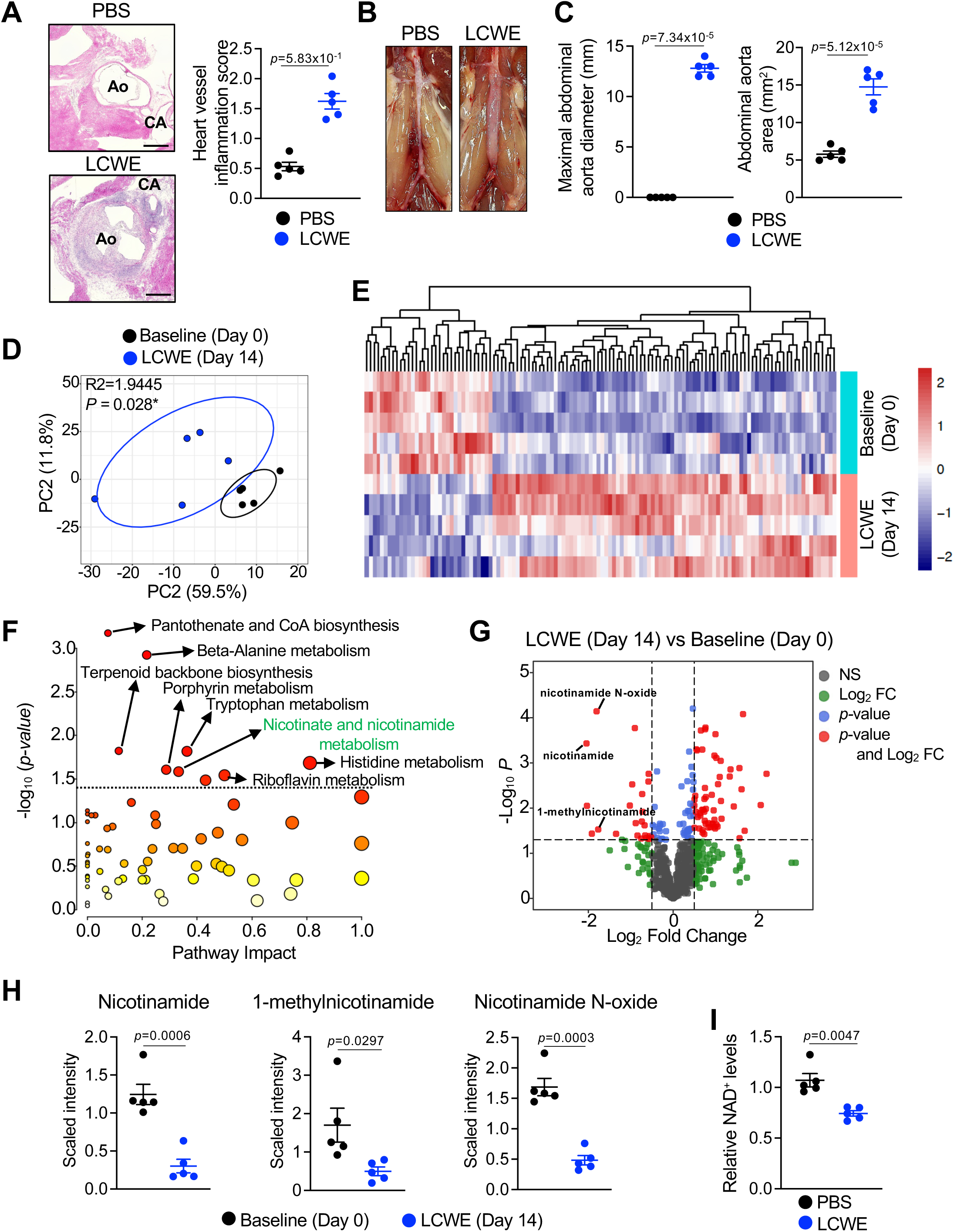
Decreased levels of NAD^+^-related metabolites in *Lactobacillus casei* cell wall extract (LCWE)-induced Kawasaki disease vasculitis. **(A)** Representative H&E-stained heart sections and heart vessel inflammation scores of WT mice injected with either PBS or LCWE at 2 weeks post-injection (n=5/group). Scale bars, 500μm. (B, C) Representative pictures of the abdominal aorta areas (B), and maximal abdominal aorta diameter and abdominal aorta area measurements (C) of WT mice injected with PBS or LCWE at 2 weeks post-injection (n=5/group). (D) Principal component analysis (PCA) plot of serum metabolome from WT mice at baseline (Day 0) or at 2 weeks post-LCWE injection (n=5/group). (E) Heatmap of significantly upregulated or downregulated (*p-value* ≤ 0.05 with paired t-test) metabolites in the serum from WT mice at baseline (Day 0) or at 2 weeks post-LCWE injection (n=5/group). (F) Metabolic pathways significantly impacted between WT mice at baseline (Day 0) and WT mice 2 weeks post-LCWE injection, determined with MetaboAnalyst. Pathways are ranked by impact scores from 0 (low impact) to 1 (strong impact), combining their threshold of significance (*p-value* ≤ 0.05) and pathway topology. Each circle denotes a pathway, and the fill color represents the significance of disturbances in that pathway from white (low significance) to red (higher significance). Dotted line indicates the threshold of significance at *p-value* ≤ 0.05 (n=5/group). (G) Volcano plot of differentially expressed metabolites in the serum of baseline (Day 0) WT mice and mice at 2 weeks post-LCWE injection. Green, log_2_ [fold change (FC)] > 0.5; blue, *p-*value ≤ 0.05; red, log_2_ [FC] > 0.5 and *p-*value ≤ 0.05) (H) Normalized scaled intensities of Nicotinamide, 1-methylnicotinamide and Nicotinamide N-oxide in the serum of WT mice at baseline (Day 0) or at 2 weeks post-LCWE injection (n=5/group). (I) NAD^+^ levels measured from serum of WT mice injected with either PBS or LCWE at 2 weeks (n=5/group). Data are presented as mean ± SEM, representative of one (C-H) or at least 2 independent experiments (A, B). Each symbol represents one individual mouse. *\*p-value* < 0.05, \*\**p-value* < 0.01, \*\*\**p-value* < 0.001, and \*\*\*\**p-value* < 0.001 by unpaired t-test (A, B, G, H). Ao, aorta, CA, coronary artery.

NAD^+^ is a metabolite central to cellular metabolism and energy production, which is also involved in essential cellular processes such as DNA repair, mitochondrial functioning, cell cycle, stress response, and cellular communication ^38^. Mammals primarily utilize NAM as a NAD^+^ precursor in the salvage pathway, with nicotinamide phosphoribosyl transferase (NAMPT) serving as the rate-limiting enzyme. NAMPT converts NAM to nicotinamide mononucleotide (NMN), which is subsequently transformed into NAD^+^ by nicotinamide/nicotinic acid mononucleotide adenylyl transferase (NMNAT) ^38^ (**Figure S1B**). SIRT1 relies on NAD^+^ for its deacetylase activity and beneficial impact on cardiovascular health. Although several poly (ADP-ribose) polymerases (PARP), cluster of differentiation (CD) 38, and CD157 also depend on NAD^+^ to exert enzymatic activity, SIRT1 accounts for a significant NAD^+^ consumption ^39^. Therefore, we next assessed SIRT1 expression in LCWE-induced cardiovascular lesions. We observed that SIRT1 protein (**Figure 2A, B**) as well as *Sirt1* mRNA levels (**Figure 2C**) were significantly decreased in heart and abdominal aortas of LCWE-injected mice compared with controls. Altogether, our results indicate that LCWE-induced vascular inflammation is associated with a downregulation in NAD^+^ metabolism and decreased SIRT1 levels.

**Figure 2.**
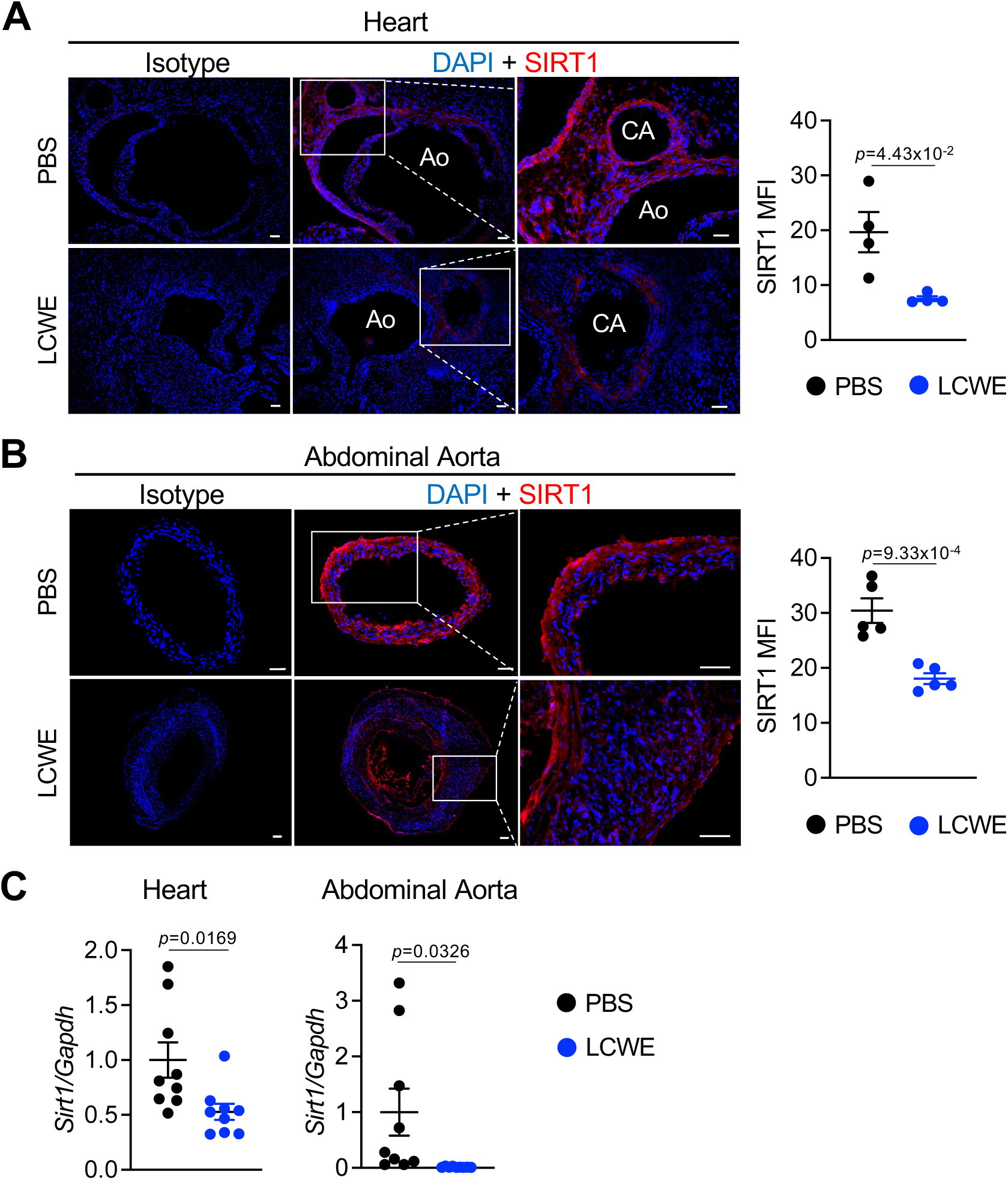
Decreased SIRT1 expression in *Lactobacillus casei* cell wall extract (LCWE)-induced cardiovascular lesions. **(A, B)** Immunofluorescent staining and mean fluorescence intensity (MFI) of SIRT1 (red) in heart (A) and abdominal aorta (B) tissues of PBS and LCWE-injected mice at 2 weeks post-LCWE injection (n=4 to 5/group). DAPI (Blue) used to stain nuclei. Scale bars, 50μm. **(C)** mRNA levels of *Sirt1* measured by qRT-PCR in heart and abdominal aorta tissues of PBS and LCWE-injected mice at 2 weeks post-LCWE injection (n=9/group). Data are presented as mean ± SEM, representative of one experiment (A, B) or pooled from 2 independent experiments (C). Each symbol represents one individual mouse. *\*p-value* < 0.05, \*\**p-value* < 0.01, \*\*\**p-value* < 0.001, and \*\*\*\**p-value* < 0.001 by unpaired t-test (B, C, D, E).

### Overexpression of SIRT1 reduces LCWE-induced cardiovascular inflammation

To determine the potential of SIRT1 to mitigate LCWE-induced KD, we used *Sirt1^super^* mice, which overexpress SIRT1 under the control of its natural promoter ^40^. Western blot (WB) analysis confirmed the increased expression of SIRT1 in heart tissues from LCWE-injected *Sirt1^super^* mice compared with LCWE-injected WT mice (**Figure S2A**). *Sirt1^super^* mice exhibited significantly less LCWE-induced heart vessel inflammation and abdominal aorta dilation when compared with WT controls (**Figure 3A-C**). LCWE injection induces an acute production of IL-1β in the peritoneal cavity ^21^. Compared with LCWE-injected WT controls, *Sirt1^super^*mice exhibited significantly reduced levels of IL-1β in their peritoneal lavage at 24 hours post-LCWE injection (**Figure 3D**).

**Figure 3.**
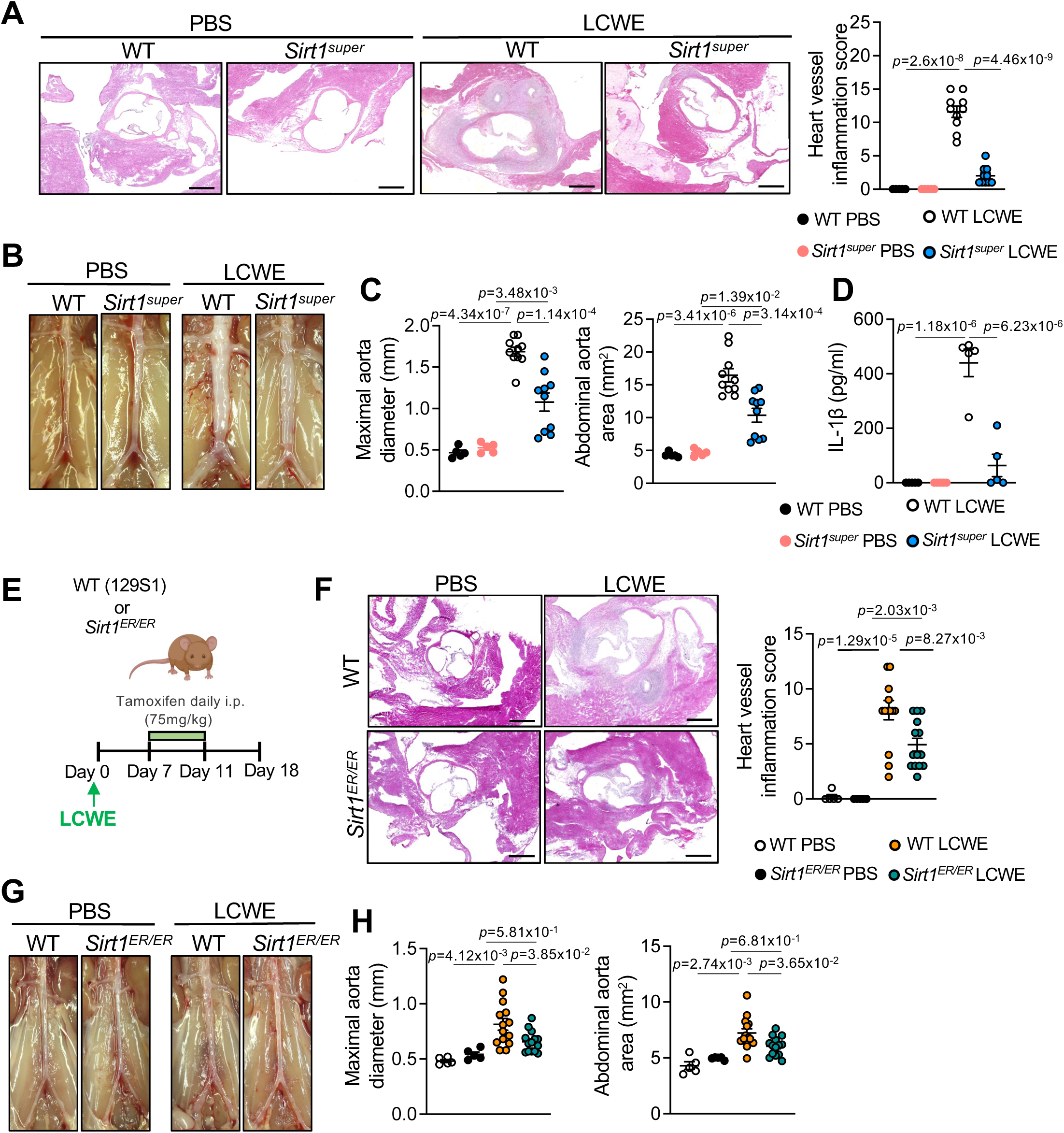
Overexpression of SIRT1 reduces *Lactobacillus casei* cell wall extract (LCWE)-induced cardiovascular inflammation. **(A)** Representative H&E-stained heart sections and heart vessel inflammation scores of PBS or LCWE-injected WT and *Sirt1^super^* mice, at 2 weeks post-injection. (n=5 to 10/group). Scale bars, 500μm. **(B, C)** Representative pictures of the abdominal aorta areas (B), maximal abdominal aorta diameter, and abdominal aorta area measurements (C) of PBS or LCWE-injected WT and *Sirt1^super^* mice, at 2 weeks post-injection. (n=5 to 10/group). **(D)** IL-1β measurements in the peritoneal lavage of WT and *Sirt1^super^* mice injected with PBS or LCWE 24 hours post-injection. **(E)** Schematic of the experimental design. **(F-H)** WT and *Sirt1^ER/ER^* mice were injected with either PBS or LCWE, and one week later, injected with tamoxifen for 5 consecutive days. Heart and abdominal aorta tissues were collected at day 18 post-LCWE injection for analysis. **(F)** Representative H&E-stained heart sections and heart vessel inflammation scores of PBS or LCWE-injected, tamoxifen-treated WT and *Sirt1^super^* mice, at day 18 post-injection. (n=5 to 14/group). Scale bars, 500μm. **(G, H)** Representative pictures of the abdominal aorta areas (G), maximal abdominal aorta diameter and abdominal aorta area measurements (H) of PBS or LCWE-injected, tamoxifen-treated WT and *Sirt1^ER/ER^* mice, at day 18 post-injection (n=5 to 14/group). Data are presented as mean ± SEM, representative of at least 2 independent experiments (A-I). Each symbol represents one individual mouse. *\*p-value* < 0.05, \*\**p-value* < 0.01, \*\*\**p-value* < 0.001, and \*\*\*\**p-value* < 0.001 by two-way ANOVA with Tukey post hoc test (A, C, D, F, H).

These results suggest that SIRT1 overexpression mitigates the development of LCWE-induced cardiovascular inflammation, which correlates with decreased LCWE-induced IL-1β production. We next determined if inducing SIRT1 expression in a therapeutic context reduced the severity of LCWE-induced KD vasculitis. We used SIRT-ER (*Sirt1^ER/ER^*) mice, in which the C-terminus of the *Sirt1* coding region is fused to the hormone-binding domain of the estrogen receptor 1α (ERα)^41^. Tamoxifen treatment of *Sirt1^ER/ER^* mice stabilizes SIRT1-ER protein, resulting in a reversible increase of its abundance and activity ^41^. Therapeutic induction of SIRT1 in *Sirt1^ER/ER^* mice significantly decreased the severity of heart vessel inflammation and abdominal aorta dilations (**Figure 3E-H**). We next treated WT and *Sirt1^ER/ER^* mice with tamoxifen for 5 days and assessed the production of IL-1β in the peritoneal lavage at 24 hours post-LCWE injection (**Figure S2B**). Compared with WT mice, induction of SIRT1-ER expression in *Sirt1^ER/ER^* mice significantly decreased acute production of IL-1β in the peritoneal cavity (**Figure S2B**). Overall, our results indicate that increasing SIRT1 levels and activity preventively or therapeutically decreases the severity of vascular inflammation and the development of cardiovascular lesions in the LCWE murine model of KD vasculitis.

### SIRT1 improves inflammation and mitochondrial function during LCWE-induced KD

To understand the molecular mechanisms underlying the protective effects of SIRT1 during LCWE-induced cardiovascular inflammation, we performed an untargeted proteomics analysis of abdominal aortas from PBS and LCWE-injected WT and *Sirt1^super^* mice (**Figure S3A**). PCA and hierarchical clustering revealed no significant differences in global protein composition of abdominal aortas between control PBS-injected WT and *Sirt1^super^* mice (**Figure 4A** and **Figure S3B**). However, significant differences in protein profiles were detected between LCWE-injected WT mice and *Sirt1^super^* mice (**Figure 4A, B,** and **Figure S3B**). We observed 1538 differentially expressed proteins (DEPs; Fold change > 2; *p-*value < 0.05) between LCWE- and PBS-injected WT mice (**Figure 4B**). This number was reduced to 511 DEPs when comparing the abdominal aortas of LCWE- and PBS-injected *Sirt1^Super^*mice (**Figure 4B**), in alignment with the finding that *Sirt1^Super^* mice develop a milder vasculitis (**Figure 3A-C**).

**Figure 4.**
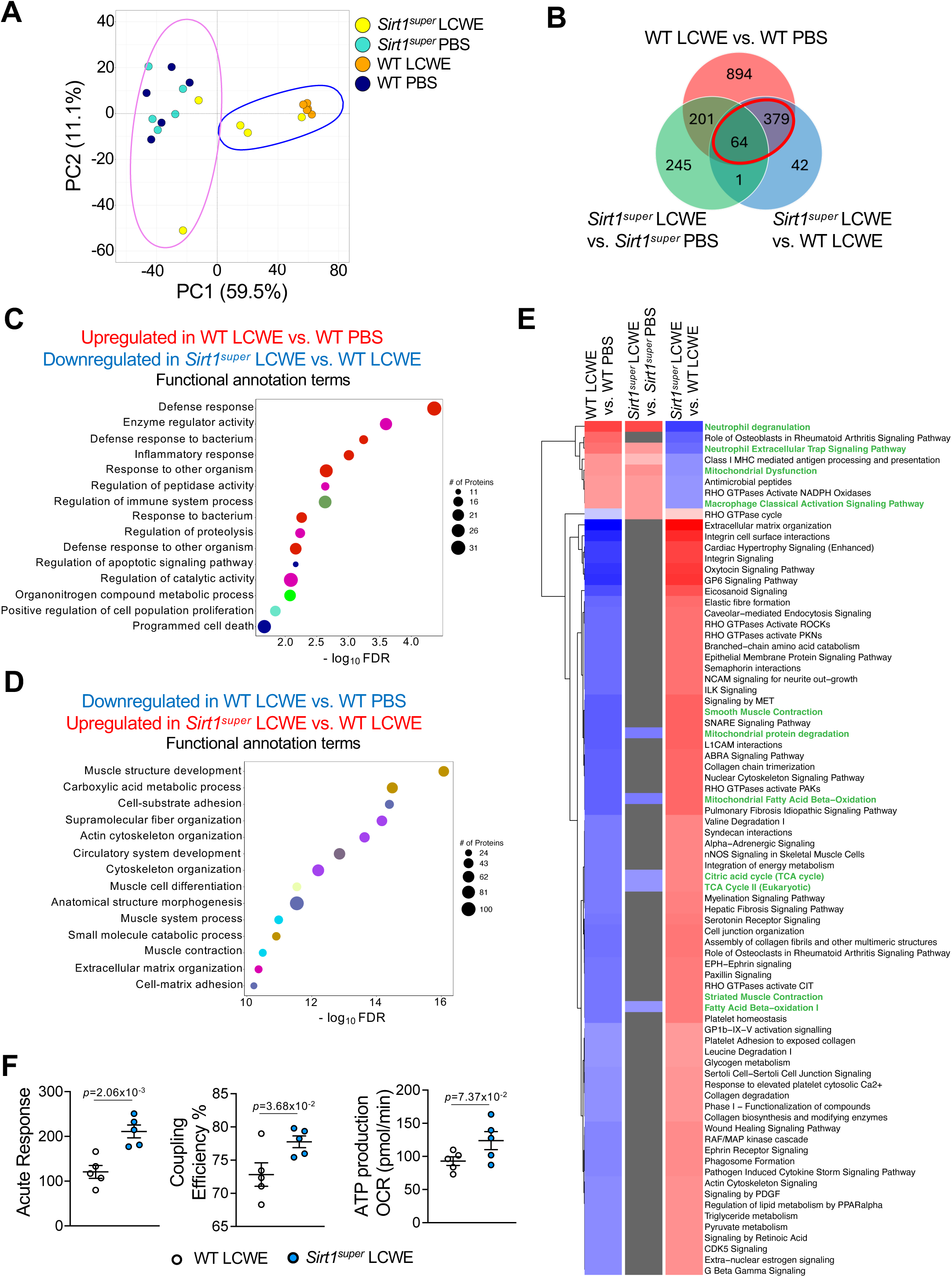
SIRT1 improves heart mitochondrial respiration, reducing *Lactobacillus casei* cell wall extract-induced cardiovascular inflammation. **(A)** Principal component (PC) analysis of the proteome from the abdominal aortas of WT and *Sirt1^super^* mice injected with either PBS or LCWE, at 2 weeks post-injection (n=5/group). **(B)** Venn diagram of differentially expressed proteins (DEPs, *p-value* <0.05; fold change [FC] >2) between the indicated groups. A set of 443 proteins associated with the development of LCWE-induced KD was identified (highlighted in red). **(C)** Selected pathways and functional annotation terms of proteins increased in LCWE-injected WT mice compared with PBS-injected WT mice and decreased in LCWE-injected *Sirt1^super^* mice compared with LCWE-injected WT mice (n=5/group). **(D)** Selected pathways and functional annotation terms of proteins decreased in LCWE-injected WT mice compared with PBS-injected WT mice and increased in LCWE-injected *Sirt1^super^* mice compared with LCWE-injected WT mice (n=5/group). **(E)** Ingenuity pathway analysis (IPA) performed on the set of DEPs associated with LCWE-induced KD highlighted in red from (B) between the indicated groups. **(F)** Mitochondrial respiration was analyzed in isolated mitochondria from heart tissues of LCWE-injected WT and *Sirt1^super^* mice. Measurements of acute response (left), coupling efficiency (middle), and ATP production (right). Data are presented as mean ± SEM, representative of one experiment. Each symbol represents one individual mouse. \**p-value* < 0.05, \*\**p-value* < 0.01, \*\*\**p-value* < 0.001, and \*\*\*\**p-value* < 0.001 by unpaired t-test (F).

We identified a set of 443 proteins associated with the development of LCWE-induced KD vasculitis based on their differential expression between LCWE and PBS-injected WT mice and differential expression between LCWE and PBS-injected *Sirt1^super^* mice (**Figure 4B**, highlighted in red). Network and pathway analysis on this set of proteins (**Figures 4C-E** and **S3C, D**) revealed an upregulation of pathways related to inflammatory and defense responses, the regulation of immune system processes and apoptosis in WT mice after LCWE-injection compared with PBS controls, which were downregulated in LCWE-injected *Sirt1^super^* mice compared with LCWE-injected WT mice (**Figure 4C** and **Figure S3C**). Furthermore, pathways related to various metabolic processes, such as fatty acid and carboxylic acid metabolism, regulation of muscle contraction, as well as circulatory system development were downregulated in LCWE-injected WT mice compared with PBS controls and upregulated in LCWE-injected *Sirt1^super^* mice compared with LCWE-injected WT mice (**Figure 4D** and **Figure S3D**).

We next performed an Ingenuity Pathway Analysis (IPA) on this set of differentially expressed proteins (**Figure 4E**). We observed a significant increase in neutrophil degranulation, neutrophil extracellular trap signaling, and macrophage activation pathways in LCWE-injected WT mice compared to PBS controls. Importantly, these pathways were significantly downregulated in LCWE-injected *Sirt1^super^* mice compared with LCWE-injected WT mice (**Figure 4E**). Furthermore, there was a significant decrease in pathways related to mitochondrial function, such as the TCA cycle, mitochondrial fatty acid oxidation, fatty acid β-oxidation and mitochondrial protein degradation in LCWE-injected WT mice compared with PBS-injected WT mice (**Figure 4E**). Strikingly, these pathways were significantly upregulated in LCWE-injected *Sirt1^super^*mice when compared with LCWE-injected WT mice (**Figure 4E**), together with a decrease in mitochondrial dysfunction, suggesting that SIRT1 promoted autophagy and mitophagy may help prevent mitochondrial dysfunction in this model ^4,42^. We also observed an increase in pathways related to smooth muscle contraction in LCWE-injected *Sirt1^super^* mice compared with LCWE-injected WT mice, suggesting a more contractile and healthier VSMC population in the abdominal aortas of these mice (**Figure 4E** and **Figure S3D**).

Given that LCWE injection, as shown in our previous work ^28^, hinders mitochondrial respiration in cardiovascular tissues, we next characterized mitochondrial respiration of freshly isolated mitochondria from heart tissues of LCWE-injected WT and *Sirt1^super^* mice. Overall, mitochondrial oxygen consumption rates (OCR) and basal glycolytic activity (extracellular acidification rate; ECAR) were similar between LCWE-injected WT and *Sirt1^super^* mice (**Figure S4A, B**). We observed a significantly higher acute response and coupling efficiency with a trend of higher ATP production (**Figure 4F**) in LCWE-injected *Sirt1^super^* mice in the same assay, suggesting that SIRT1 overexpression improves mitochondrial respiration in heart tissues. However, no differences in levels of heart tissue mtDNA were observed between LCWE-injected WT and *Sirt1^super^* mice at 2 weeks post-LCWE injection, indicating that the enhanced respiration in *Sirt1^super^*mice was not related to a change in mitochondrial mass (**Figure S4C).** These results suggest that SIRT1 counteracts the inflammatory response and promotes mitochondrial function during LCWE-induced KD vasculitis.

### SIRT1 reduces the accumulation of ROS in cardiovascular lesions and improves autophagy/mitophagy during LCWE-induced KD

Accumulation of dysfunctional mitochondria due to impaired autophagy leads to increased production of ROS, intensifying the oxidative stress response in cardiovascular diseases ^43,44^. We previously showed that LCWE-induced KD vasculitis is linked to impaired clearance of dysfunctional mitochondria and increased ROS accumulation in the cardiovascular lesions of LCWE-injected WT mice compared with PBS control mice ^28^. Therefore, we next assessed the expression of genes related to autophagy, ROS metabolism, and mitochondrial respiration in two publicly available gene expression datasets generated from the whole blood of acute KD and healthy control (HC) patients (GSE68004 and GSE73461; **Figure S5**). The expression of various genes involved in scavenging ROS (*SOD1*, *PRDX1*, *UCP2*) was decreased in acute KD patients compared with HC in both datasets (**Figure S5A**). We also detected downregulation in the expression of *BNIP3*, a mitophagy receptor, as well as downregulation of genes known to modulate autophagy pathways by blocking mTOR, such as *DDIT4* and *SESN1* (**Figure S5A**, GSE68004). Similarly, multiple genes encoding electron transport chain complex subunits were also downregulated during acute KD compared with HC in the same datasets (**Figure S5B**). These results indicate that pathways linked to autophagy, ROS scavenging, and oxidative phosphorylation are decreased in patients with KD during the acute phase of the disease.

To determine if SIRT1 overexpression impacts oxidative stress, we next quantified local ROS levels in hearts and abdominal aortas of WT and *Sirt1^super^* mice by dihydroethidium (DHE) staining. While LCWE injection significantly increased ROS production in both hearts and abdominal aortas of WT mice, it was strongly decreased in LCWE-injected *Sirt1^super^* mice (**Figure 5A** and **Figure S6A**). To assess whether SIRT1 overexpression improves impaired autophagy/mitophagy, ROS accumulation, and mitochondrial respiration during LCWE-induced KD, we analyzed protein expression in the abdominal aortas from PBS or LCWE-injected WT and *Sirt1^super^* mice (**Figure 5B**). While the levels of proteins related to autophagy/mitophagy/ROS, fatty acid oxidation, and OXPHOS appear identical in the abdominal aorta of PBS-injected control WT and *Sirt1^super^* mice, most of these proteins were significantly reduced in the inflamed abdominal aorta of LCWE-injected WT mice (**Figure 5B**). However, this effect was mitigated in the abdominal aorta of LCWE-injected *Sirt1^super^* mice, which develop a less severe LCWE-induced KD (**Figure 5B**). In particular, the expression of microtubule-associated protein 1a/1b light chain 3a (Map1lc3a), which is involved in autophagosome formation and selection of autophagic cargo, was reduced by LCWE injection in WT mice, indicating impaired mitophagy (**Figure 5B**). However, overexpression of SIRT1 in *Sirt1^super^* mice attenuated this LCWE-induced alteration in the expression of Map1lc3a, nearly normalizing the level of this protein to that observed in control PBS-injected WT and *Sirt1^super^* mice, suggesting functional mitophagy (**Figure 5B**). Furthermore, the levels of mitochondrial uncoupling protein 1 (Ucp1) were significantly higher in LCWE-injected *Sirt1^super^* mice compared with LCWE-injected WT mice, hinting towards a more efficient machinery of ROS scavenging (**Figure 5B**). Various proteins involved in fatty acid oxidation and oxidative phosphorylation were also upregulated in LCWE-injected *Sirt1^super^* mice (**Figure 5B**), further supporting our previous results that SIRT1 overexpression promotes efficient mitochondrial oxidative phosphorylation.

**Figure 5.**
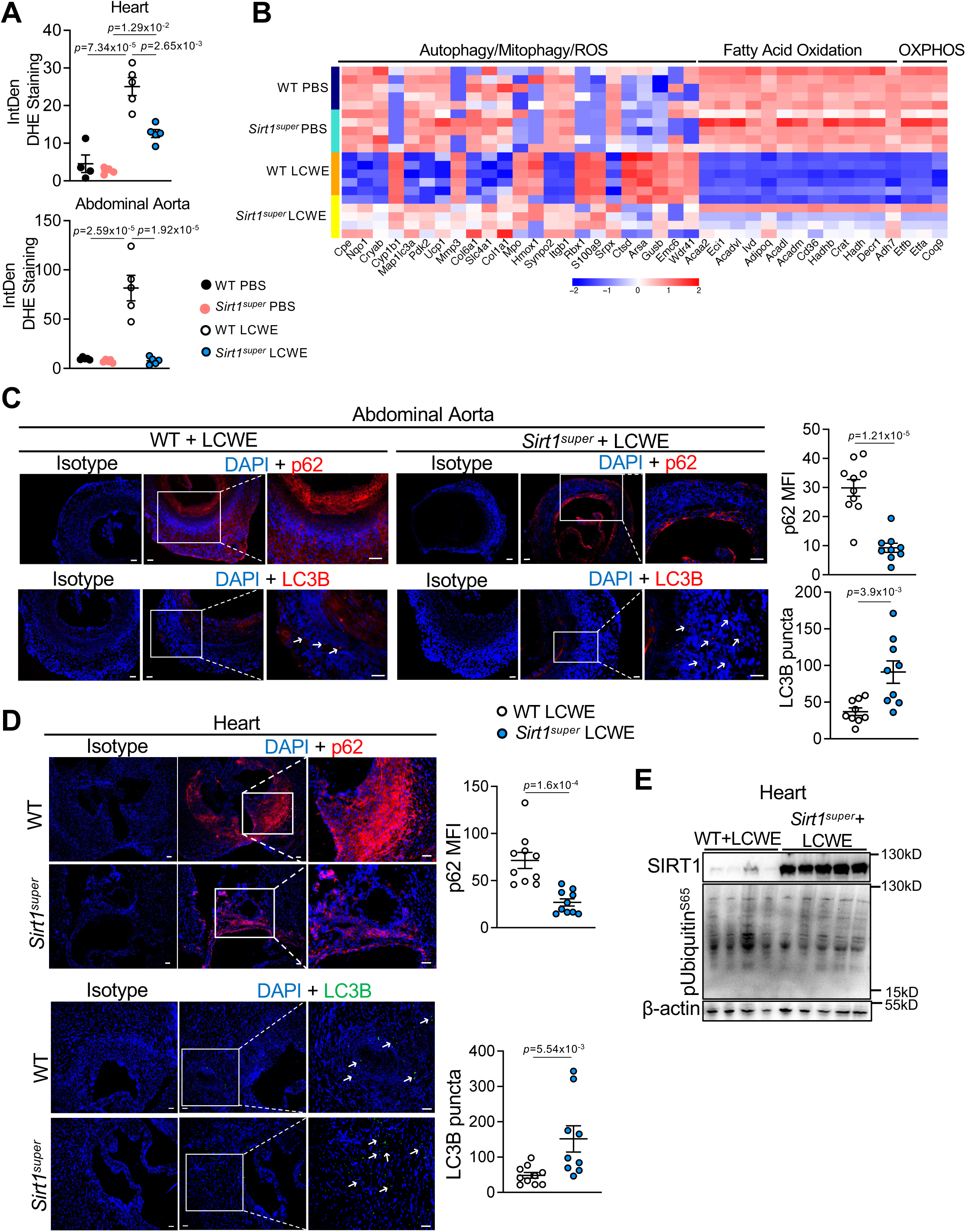
Overexpression of SIRT1 reduces ROS accumulation and promotes autophagy/mitophagy during *Lactobacillus casei* cell wall extract (LCWE)-induced KD. **(A)** Quantification of dihydroethidium (DHE) staining of heart (top) and abdominal aorta (bottom) tissues from WT and *Sirt1^super^* mice injected with either PBS or LCWE at 2 weeks post-injection (n*=*4 to 5/group). **(B)** Heatmap of differentially expressed proteins related to autophagy/ROS, fatty acid oxidation, and oxidative phosphorylation pathways between abdominal aortas of PBS and LCWE-injected WT and *Sirt1^super^* mice at 2 weeks post-injection (n=5/group). **(C)** Immunofluorescent staining and quantification of p62 (red) and LC3B puncta (red) in abdominal aorta tissues from PBS or LCWE-injected WT and *Sirt1^super^* mice at 2 weeks post-injection (n=9,10/group). DAPI (blue) is used to stain nuclei. Scale bars, 50μm. **(D)** Immunofluorescent staining and quantification of p62 (red) and LC3B puncta (green) from hearts of PBS or LCWE-injected WT and *Sirt1^super^* mice at 2 weeks post-injection (n=9,10/group). DAPI (blue) is used to stain nuclei. Scale bars, 50μm. **(E)** Western blot analysis of SIRT1, pUbiquitin^S65^ and β-actin in protein lysates from heart tissues from WT and *Sirt1^super^* mice injected with LCWE at 2 weeks post-injection (n=4, 5/group). Data are presented as mean ± SEM. Pooled from 2 experiments (A, C, D, E), representative of one experiment (B). Each symbol represents one individual mouse. *\*p-value* < 0.05, \*\**p-value* < 0.01, \*\*\**p-value* < 0.001, and \*\*\*\**p-value* < 0.001 by two-way ANOVA with Tukey post hoc test (A) or unpaired t-test (C, D). MFI, mean fluorescence intensity.

We next determined the protein levels of p62, a protein that binds to ubiquitinated targets and is rapidly degraded during active mitophagy ^44^, and LC3B, which forms visible puncta during autophagosome formation, allowing assessment of autophagosome numbers by immunofluorescent staining in cardiovascular tissues (**Figure 5C, D**). We observed a significant decrease in p62 levels, whereas LC3B puncta formation was significantly increased in both abdominal aortas and heart tissues of LCWE-injected *Sirt1^super^* mice compared with LCWE-injected control WT mice, suggesting intact mitophagy when SIRT1 is overexpressed (**Figure 5C, D**). We have previously shown that in WT mice, LCWE-induced vasculitis correlates with increased heart levels of phospho-ubiquitin Ser65 (pUbiquitin^S65^) ^28^, a Pink1/Parkin mitophagy pathway-specific posttranslational modification of ubiquitin ^45^. Overexpression of SIRT1 counteracts impaired mitophagy in LCWE-injected mice, as evidenced by a significant decrease of pUbiquitin^S65^ levels in heart tissues of LCWE-injected *Sirt1^super^* mice compared with WT control mice, indicating ubiquitination followed by degradation of target mitochondrial cargo (**Figure 5E** and **Figure S6B**). To further understand the regulation of autophagy/mitophagy by SIRT1, we performed an *in vivo* autophagic flux assay in LCWE-injected WT and *Sirt1^super^* mice ^28,46^. Mice were injected with LCWE, and 2 weeks later, a subset received a single dose of chloroquine (CQ) to inhibit autophagosome-lysosome fusion (**Figure S6C**). Abdominal aortas were harvested 4 hours after CQ, and accumulation of LC3-II and p62, corresponding to autophagic flux, was assessed by WB analysis (**Figure S6D, E**). As previously published, LC3-II accumulation was impaired in abdominal aortas of LCWE-injected WT mice treated with CQ, confirming defective autophagy during LCWE-induced KD (**Figure S6D, E**) ^28^. In contrast, the increased accumulation of p62 and LC3-II in abdominal aorta tissues of LCWE-injected *Sirt1^super^* mice following CQ treatment suggests that SIRT1 overexpression promotes induction of autophagy during LCWE-induced KD (**Figure S6D, E**). Overall, our results indicate that overexpression of SIRT1 promotes the clearance of damaged mitochondria and decreases ROS accumulation during LCWE-induced cardiovascular inflammation.

### SIRT1 overexpression boosts contractile gene expression and autophagy/mitophagy in smooth muscle cells and suppresses macrophage IL-1β production

During LCWE-induced KD, VSMCs undergo a phenotypic switch, becoming synthetic VSMCs or type II VSMCs ^21^. This process is driven by IL-1 signaling and results in increased expression of chemokines and fibroblastic markers, decreased expression of contractile proteins, and enhanced proliferative capacity ^21^. Confirming these observations, we detected a decrease in pathways related to smooth muscle contraction in LCWE-injected WT mice in the IPA of our proteomics data, which was reversed in LCWE-injected *Sirt1^super^*mice (**Figure 4E**). More specifically, we observed reduced expression of proteins related to contractile functions of VSMCs (Tagln, Myh11, Ppp1r14a, Ppp1r12b, Myl9, Mylk, Itga8, Myl6, Gucy1b1, Lmod1, Rock1, Adcy5, Itga9, Ncam1; Type I VSMC marker), as well as increased expression of matrix metalloproteinase (Mmp) 3 and 14 (a Type II VSMC marker) in the abdominal aortas of LCWE-injected WT mice (**Figure 6A**). All these alterations in protein expression were strongly attenuated in LCWE-injected *Sirt1^super^*mice, indicating that overexpression of SIRT1 mitigated the LCWE-induced VSMC phenotypic switch (**Figure 6A**). Immunofluorescent staining showed increased expression of Lumican, a type II VSMC protein, and decreased expression of the type I VSMC-related protein αsmooth muscle actin (αSMA) in both heart and abdominal aorta tissues of LCWE-injected WT mice, further confirming the VSMCs’ phenotypic switch (**Figure S7A-D**). VSMCs surrounding the aorta and coronary arteries significantly expressed higher levels of αSMA, and lower levels of Lumican in cardiovascular tissues from LCWE-injected *Sirt1^super^*mice (**Figure S7A-D**). Furthermore, mRNA levels of *Mmp9* (type II VSMC marker) were significantly lower, while the expression of *Myh11* (type I VSMC marker) was significantly higher in primary VSMCs isolated from abdominal aortas of *Sirt1^super^* mice stimulated with LCWE or recombinant IL-1β (rIL-1β) *in vitro* (**Figure S7E**). Altogether, our results indicate that SIRT1 overexpression promotes efficient mitochondrial respiration (**Figure 4** and **Figure S4**) and counteracts the synthetic phenotypic switch of VSMCs towards type II VSMCs during LCWE-induced KD (**Figure 6** and **Figure S7**). To further explore the regulation of autophagy/mitophagy by SIRT1 *in vitro*, we next used primary VSMCs from WT mice. We stimulated them with LCWE in the presence or absence of SRT1720, a synthetic SIRT1 activator ^47,48^, or EX527, a synthetic SIRT1 inhibitor ^49,50^, and assessed their expression of autophagy/mitophagy-related proteins by WB analysis. Treatment of LCWE-stimulated VSMCs with the SIRT1 activator SRT1720 increased autophagy, as indicated by increased LC3-II levels and decreased pS6^S235/236^ levels, but this effect was not observed when VSMCs were treated with the SIRT1 inhibitor EX527 (**Figure 6B** and **Figure S8A**). SRT1720 treatment of LCWE-stimulated VSMCs did not alter the levels of phosphorylated Unc-51-like kinase 1 (ULK1) and AMP-activated protein kinase (AMPK), two proteins that promote autophagy ^51^ (**Figure 6B** and **Figure S8A**). These results indicate that SIRT1 promotes autophagy/mitophagy, potentially by counteracting mTOR’s suppressive activity.

**Figure 6.**
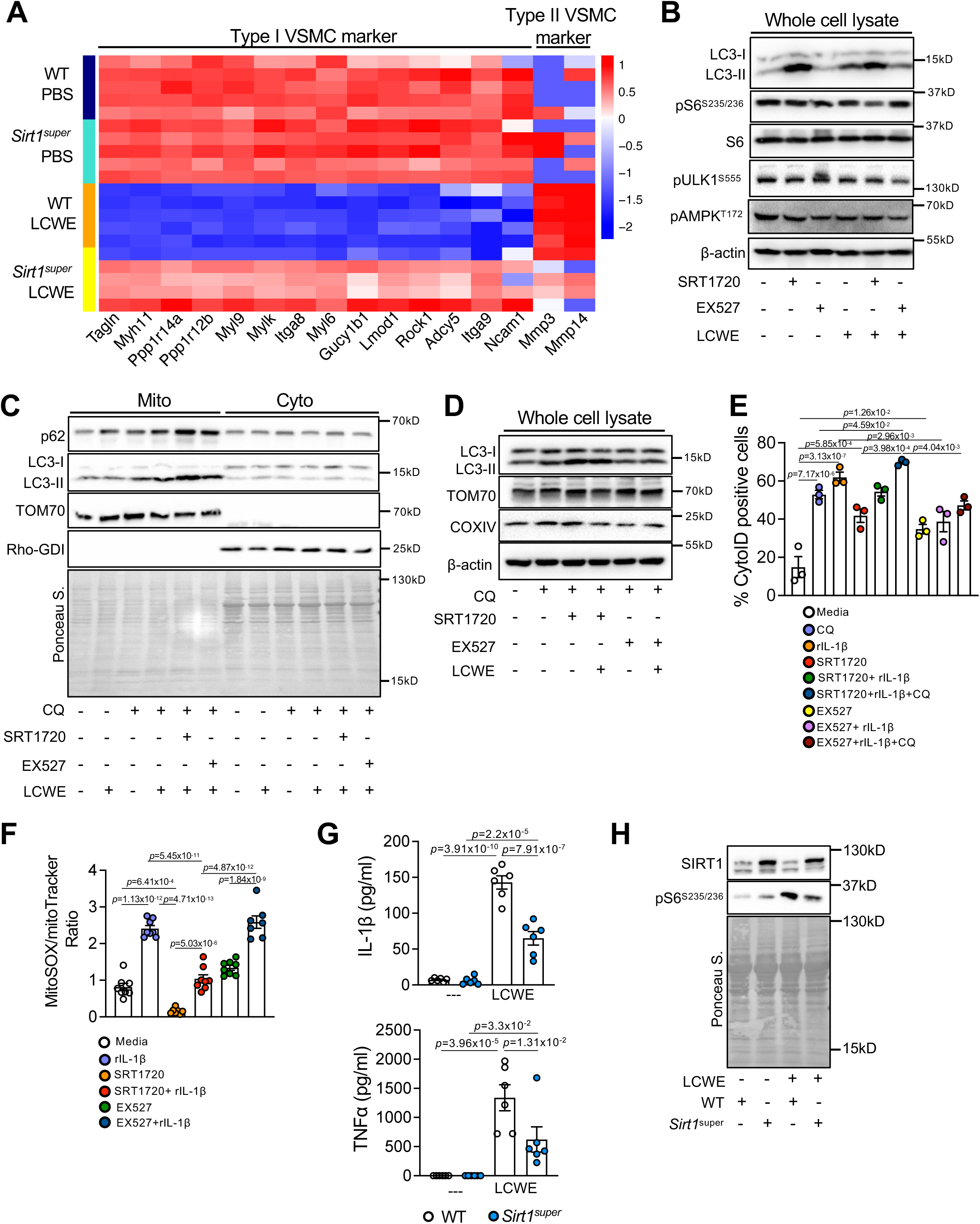
SIRT1 promotes autophagy/mitophagy in vascular smooth muscle cells (VSMCs), blocking their phenotypic switch, and suppresses IL-1β secretion from macrophages. **(A)** Heatmap of differentially expressed proteins (DEPs) related to VSMCs type I and synthetic VSMCs type II in the abdominal aortas of PBS and LCWE-injected WT and *Sirt1^super^* mice at 2 weeks post-injection (n=5/group). **(B)** Primary coronary artery SMCs were treated with SRT1720 (5μM) or EX527 (5μM) alone or in combination with LCWE (60μg/ml, 24 hours). Representative Western blots of LC3-I/II, pS6^S235/236^, S6, pULK1^S555^, pAMPK^T172^ and β-actin from whole cell lysate (n=4/group). **(C)** Representative Western blots of p62, LC3-I/II, TOM70 and Rho-GDI from Mitochondrial (Mito) and cytosolic (Cyto) fractions of primary coronary artery SMCs treated with SRT1720 (5μM) or EX527 (5μM) alone or in combination with LCWE (60μg/ml, 24 hours) and CQ (5μM, last 4 hours). Ponceau S. was used as loading control. **(D)** Representative Western blots of LC3-I/II, TOM70, COXIV and β-actin from whole cell lysate of primary coronary artery SMCs treated with SRT1720 (5μM) or EX527 (5μM) alone or in combination with LCWE (60μg/ml, 24 hours) and CQ (5μM, last 4 hours). **(E)** Quantification of CytoID staining in WT primary coronary artery SMCs treated with SRT1720 (5μM, 24 hours) or EX527 (5μM, 24 hours) alone or in combination with rIL-1β (10ng/ml, 24 hours) and CQ (5μM, last 4 hours) (n=3/group). **(F)** Quantification of mitoSOX/mitoTracker ratio from WT Primary coronary artery SMCs treated with SRT1720 (5μM, 24 hours) or EX527 (5μM, 24 hours) alone or in combination with rIL-1β (10ng/ml, 24 hours) (n=8/group). **(G-H)** Bone marrow-derived macrophages (BMDMs) differentiated from WT and *Sirt1^super^* mice treated with LCWE (60μg/ml, 24 hours). **(G)** IL-1β and TNFα levels were measured from the supernatants by ELISA (n=6/group). **(H)** Representative Western blots for SIRT1 and pS6^S235/236^ from whole cell lysates (n=4/group). Ponceau S. was used as loading control. Data are presented as mean ± SEM. Representative of 2 experiments (B, C, D, E, F, G, H), representative of one experiment (A). Each symbol represents one individual mouse. *\*p-value* < 0.05, \*\**p-value* < 0.01, \*\*\**p-value* < 0.001, and \*\*\*\**p-value* < 0.001 by one-way ANOVA (E, F) or two-way ANOVA with Tukey post hoc test (H). rIL-1β, recombinant IL-1β.

We next enriched the mitochondrial fraction of LCWE-stimulated and SRT1720 or EX527-treated VSMCs, to measure autophagic flux by WB analysis ^46^ (**Figure 6C**). The accumulation of p62 and LC3-II in the mitochondrial fraction was significantly higher in cells treated with SRT1720 (**Figure 6C** and **Figure S8B**), indicating that boosting SIRT1 activity promotes autophagic flux *in vitro* after LCWE stimulation. Increased LC3-II levels in whole cell lysate paralleled the increase observed in the mitochondrial fraction, further confirming a higher autophagic flux when SIRT1 is activated (**Figure 6D** and **Figure S8C**). Notably, the levels of the mitochondrial protein translocase of outer mitochondrial membrane 70 (TOM70) and cytochrome c oxidase subunit 4 (COX IV) remained unchanged (**Figure 6D** and **Figure S8C**), in line with our finding that SIRT1 does not impact mitochondrial mass in the LCWE-induced KD murine model.

We previously reported that in LCWE-induced cardiovascular lesions, VSMCs respond to IL-1β, which triggers their synthetic phenotypic switch ^21^. IL-1β has also been associated with the induction of autophagic flux and mitochondrial damage ^52,53^. Therefore, we next assessed autophagic flux *in vitro* in primary VSMCs stimulated with either CQ, IL-1β, or a combination of CQ and IL-1β, using CytoID dye, which labels autophagic compartments ^54^. CQ and IL-1β stimulation increased autophagic flux in VSMCs, as demonstrated by higher frequencies of CytoID^+^ cells. Treatment of VSMCs with the SIRT1 activator SRT1720 boosted CytoID^+^ cell numbers even more during IL-1β-CQ-mediated induction of autophagic flux, whereas SIRT1 inhibitor EX527 reduced these numbers (**Figure 6E** and **Figure S9A**). Furthermore, stimulation of primary VSMCs with rIL-1β promoted accumulation of mitochondrial ROS (**Figure 6F** and **Figure S9B**). Activating SIRT1 in these cells with SRT1720 significantly decreased the accumulation of mitochondrial ROS (**Figure 6F** and **Figure S9B**), consistent with our *in vivo* results (**Figure 5A** and **S5A**). Overall, these results suggest that SIRT1 activation enhances autophagic flux and reduces ROS formation in VSMCs *in vitro*.

Since in LCWE-induced cardiovascular lesions, the major sources of IL-1β are monocytes and macrophages ^16,21,55^, we next sought to determine if SIRT1 overexpression impacts their production of IL-1β. Compared with LCWE-stimulated WT bone marrow-derived macrophages (BMDMs), the production of IL-1β and tumor necrosis factor alpha (TNF-α) was decreased in LCWE BMDMs generated from *Sirt1^super^* mice (**Figure 6G**). Furthermore, we also observed a significant decrease in pS6^S235/236^ levels in BMDMs from *Sirt1^super^* mice, confirming SIRT1’s inhibitory role on mTOR activity (**Figure 6H** and **Figure S9C**). Overall, our results indicate that SIRT1 decreases NLRP3 inflammasome activation and IL-1β production by macrophages, enhances VSMCs autophagic flux, and reduces ROS production *in vitro* and *in vivo*

### Specific deletion of Sirt1 in vascular smooth muscle cells or macrophages increases the severity of LCWE-induced KD vasculitis

To determine the impact of SIRT1 on VSMC autophagy *in vivo*, we generated mice with a tamoxifen-inducible specific deletion of *Sirt1* in VSMCs by crossing *Myh1^Cre/ERT2^* mice with *Sirt1^fl/fl^*mice. *Myh11^Cre/ERT2^Sirt1^Δ/Δ^* and *Sirt1^fl/fl^* littermate controls were fed a tamoxifen diet for 2 weeks to induce *Sirt1* deletion, and after a 4-week rest period, mice were injected with LCWE to induce vasculitis (**Figure S10A**). SIRT1 expression was reduced in the abdominal aortas of tamoxifen-treated *Myh11^Cre/ERT2^Sirt1^Δ/Δ^* when compared with *Sirt1^fl/fl^*littermate control mice (**Figure S10B**). Furthermore, *in vitro* 4-hydroxytamoxifen (4OHT) treatment of primary VSMCs resulted in *Sirt1* deletion in VSMCs from *Myh11^Cre/ERT2^Sirt1^Δ/Δ^* mice and not VSMCs from *Sirt1^fl/fl^* littermate controls (**Figure S10C**). Deletion of *Sirt1* in VSMCs of *Myh11^Cre/ERT2^Sirt1^Δ/Δ^* mice resulted in the development of more severe LCWE-induced cardiovascular inflammation when compared with littermate *Sirt1^fl/fl^* controls (**Figure 7A-C**). However, production of IL-1β in the peritoneal cavity at 24 post-LCWE injection was not affected, as monocytes and macrophages are the primary source of IL-1β production (**Figure 7D**) ^21,22,56^. Furthermore, consistent with our findings that SIRT1 overexpression promotes the clearance of dysfunctional mitochondria (**Figures 5 and 6**), deletion of *Sirt1* in VSMCs resulted in accumulation of p62 and a decreased LC3-II/I ratio after stimulation with IL-1β, suggesting impaired mitophagy/autophagy (**Figure S10C**). These results suggest that lack of *Sirt1* in VSCMs exacerbates LCWE-induced KD by interfering with autophagy/mitophagy. As monocytes and macrophages are the main source of bioactive IL-1β in LCWE-induced KD vasculitis ^21,22,56^, we next sought to determine the impact of *Sirt1* deletion in myeloid cells during LCWE-induced KD by crossing *Csf1r^Cre^* with *Sirt1^fl/fl^*mice (**Figure S10D**). Deletion of *Sirt1* in myeloid cells worsened the severity of LCWE-induced KD, as demonstrated by increased heart inflammation and abdominal aorta dilations in LCWE-injected *Csf1r^Cre^Sirt1^Δ/Δ^*mice (**Figure 7E-G**). Furthermore, these mice also had a significant induction of peritoneal IL-1β production at 24 hours post-LCWE injection (**Figure 7H**). *In vitro*, compared with BMDMs differentiated from *Sirt1^fl/fl^* littermate control mice, BMDMs from *Csf1r^Cre^Sirt1^Δ/Δ^* mice showed significantly higher levels of secreted IL-1β and TNFα after LCWE stimulation (**Figure S10E**). *Sirt1* deletion in BMDMs resulted in increased M1 macrophage polarization after lipopolysaccharide (LPS) and interferon gamma (IFN*γ*) stimulation, as demonstrated by higher mRNA expression of inducible nitric oxide synthase (*Nos2*) and *Tnfa* (**Figure S10F**). Furthermore, IL-4-induced M2 polarization was reduced in BMDMs from *Csf1r^Cre^Sirt1^Δ/Δ^*mice, with significantly less expression of found in inflammatory zone 1 (*Fizz1*) and chitinase-like 3 (*Ym1* or *Chil3*) (**Figure S10F**). LCWE-induced KD vasculitis leads to the infiltration of immune cells, including F4/80^+^ cells with Caspase-1 activity (fluorochrome-labeled inhibitors of caspases [FLICA]^+^), into cardiovascular tissues ^21,22,56,57^. We next assessed the number of infiltrating F4/80^+^ cells that are FLICA and NLRP3-positive in cardiovascular tissues of control or *Csf1r^Cre^Sirt1^Δ/Δ^*mice injected with PBS or LCWE (**Figure 7I, J,** and **Figure S11A, B**). Loss of *Sirt1* in myeloid cells was linked to higher numbers of F4/80, FLICA, and NLRP3-positive cells infiltrating cardiovascular tissues after LCWE-injection, compared with *Sirt1^fl/fl^* littermate control mice (**Figure 7I, J,** and **Figure S11A, B**). Overall, these results indicate that loss of SIRT1 in VSMCs and macrophages exacerbates IL-1β and TNFα production, drives impaired autophagy/mitophagy and favors proinflammatory M1-like macrophage polarization, thus promoting cardiovascular lesions during LCWE-induced KD.

**Figure 7.**
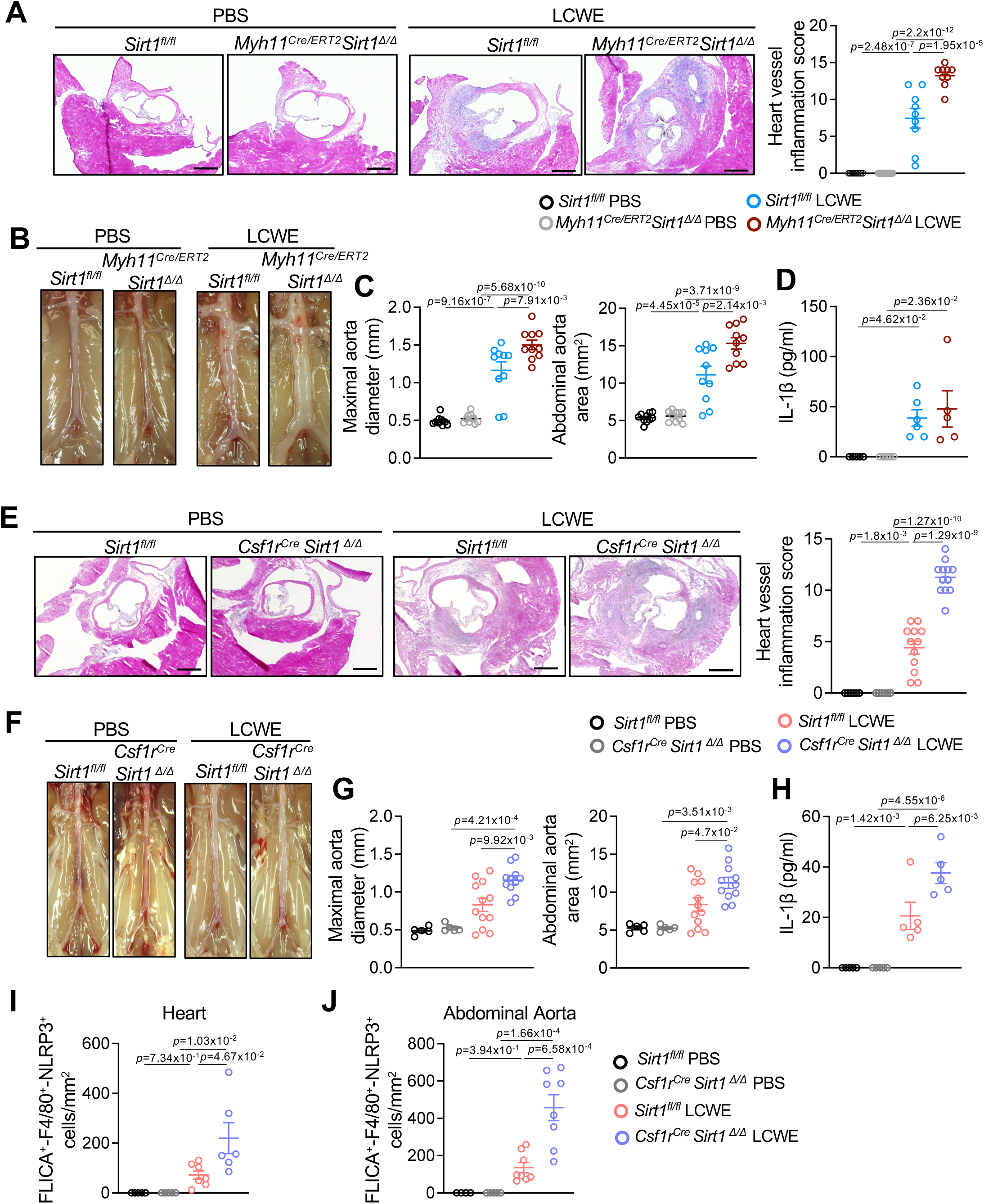
Specific deletion of *Sirt1* in vascular smooth muscle cells or myeloid cells increases the severity of *Lactobacillus casei* cell wall extract (LCWE)-induced KD vasculitis. **(A)** Representative H&E-stained heart sections and heart vessel inflammation scores of *Sirt1^fl/fl^* and *Myh11^Cre/ERT2^Sirt1^Δ/Δ^* mice were injected with PBS or LCWE and analyzed at 2 weeks post-injection (n=9 to 10/group). Scale bars, 500μm. **(B, C)** Representative pictures of the abdominal aorta areas (B), maximal abdominal aorta diameter and abdominal aorta area (C) measurements of *Sirt1^fl/fl^* and *Myh11^Cre/ERT2^Sirt1^Δ/Δ^* mice were injected with PBS or LCWE and analyzed at 2 weeks post-injection (n=9 to 10/group). **(D)** Levels of IL-1β in the peritoneal lavage of *Sirt1^fl/fl^* and *Myh11^Cre/ERT2^Sirt1^Δ/Δ^* mice injected with PBS or LCWE 24 hours post-injection. (E-G) *Sirt1^fl/fl^* and *Csf1r^Cre^Sirt1^Δ/Δ^* mice were injected with PBS or LCWE and analyzed at 2 weeks post-injection (n=5 to 11-13/group). **(E)** Representative H&E-stained heart sections and heart vessel inflammation scores of *Sirt1^fl/fl^* and *Csf1r^Cre^Sirt1^Δ/Δ^* mice were injected with PBS or LCWE and analyzed at 2 weeks post-injection (n=5 to 11-12/group). Scale bars, 500μm. **(F, G)** Representative pictures of the abdominal aorta areas (F), maximal abdominal aorta diameter, and abdominal aorta area measurements (G) of *Sirt1^fl/fl^* and *Csf1r^Cre^Sirt1^Δ/Δ^* mice were injected with PBS or LCWE and analyzed at 2 weeks post-injection (n=5 to 11-12/group). **(H)** Levels of IL-1β in the peritoneal lavage of *Sirt1^fl/fl^* and *Csf1r^Cre^Sirt1^Δ/Δ^* mice injected with PBS or LCWE 24 hours post-injection (n=5/group). **(I)** Quantification of FLICA^+^F4/80^+^NLRP3^+^ cell numbers in heart tissues from PBS or LCWE injected *Sirt1^fl/fl^* and *Csf1r^Cre^Sirt1^Δ/Δ^* mice at 2 weeks post-injection (n=5 to 7/group). **(J)** Quantification of FLICA^+^ F4/80^+^NLRP3^+^ cell numbers in abdominal aortas from PBS or LCWE injected *Sirt1^fl/fl^* and *Csf1r^Cre^Sirt1^Δ/Δ^* mice at 2 weeks post-injection (n=4 to 5-8/group). Data are presented as mean ± SEM. Pooled from 2 experiments (A-J). Each symbol represents one individual mouse. *\*p-value* < 0.05, ** *p-value* < 0.01, *** *p-value* < 0.001, and **** *p-value* < 0.001 by two-way ANOVA with Tukey post hoc test (A, C, D, E, G, H, I, J). FLICA, Fluorochrome Labeled Inhibitors of Caspases.

### Boosting SIRT1 activity and NAD^+^ levels decreases the severity of LCWE-induced KD vasculitis

We next determined if activating SIRT1 *in vivo* reduces the severity of LCWE-induced KD vasculitis. We first supplemented WT mice with resveratrol (RES), a compound known for its capacity to activate SIRT1 ^48,58,59^. WT mice received RES daily by oral gavage starting the day before LCWE injection, until the experimental endpoint (**Figure S12A**). RES treatment significantly decreased the severity of LCWE-induced heart vessel inflammation and abdominal dilations (**Figure S12B-D**). However, RES can also interact with other Sirtuins and influence a wide range of anti-inflammatory molecular pathways ^60–62^. Therefore, we next assessed the impact of SRT1720 on LCWE-induced KD vasculitis *in vivo*, a synthetic SIRT1 activator ^47,48^. Compared to LCWE-injected mice, daily treatment with SRT1720, starting the day before LCWE injection, significantly decreased the severity of LCWE-induced cardiovascular lesions (**Figure 8A-C** and **Figure S12E**). Since we observed decreased levels of NAD^+^-related metabolites and NAD^+^ activity in LCWE-injected WT mice developing KD vasculitis (**Figure 1**) and SIRT1 activity is regulated by NAD^+^ levels ^63,64^, we reasoned that boosting NAD^+^ production by supplementing with its precursors, β-nicotinamide mononucleotide (β-NMN) or nicotinamide riboside (NR), may reduce the severity of LCWE-induced KD. We supplemented mice with β-NMN or NR, starting one day before LCWE injection and continuing until the experimental endpoint (**Figure S12F, G**). Prophylactic treatment of LCWE-injected mice with β-NMN (**Figure 8D-F**) or NR (**Figure 8G-I**) decreased the severity of LCWE-induced heart vessel inflammation and the development of abdominal aorta dilations. This reduction in LCWE-induced vascular inflammation was associated with increased NAD^+^ levels (**Figure S12H, I**). We next assessed if providing NAD^+^ precursors therapeutically, after LCWE injection, would also impact the development of LCWE-induced cardiovascular lesions. Mice were first injected with LCWE and then supplemented with β-NMN or NR starting on day 3 post-LCWE injection (**Figure S13A**). Compared with LCWE-injected WT mice, we observed a therapeutic effect and a significant reduction in heart vessel inflammation when the mice were treated with β-NMN on day 3 post-LCWE injection (**Figure S13B-D**), while NR treatment decreased both heart vessel inflammation and the development of abdominal aorta aneurysms after LCWE injection (**Figure S13E-G**). Altogether, our data indicate that increasing NAD^+^ levels or SIRT1 activity counteracts the severity of LCWE-induced KD vasculitis, highlighting the NAD^+^/SIRT1 axis as a promising therapeutic target to mitigate the development and progression of KD.

**Figure 8.**
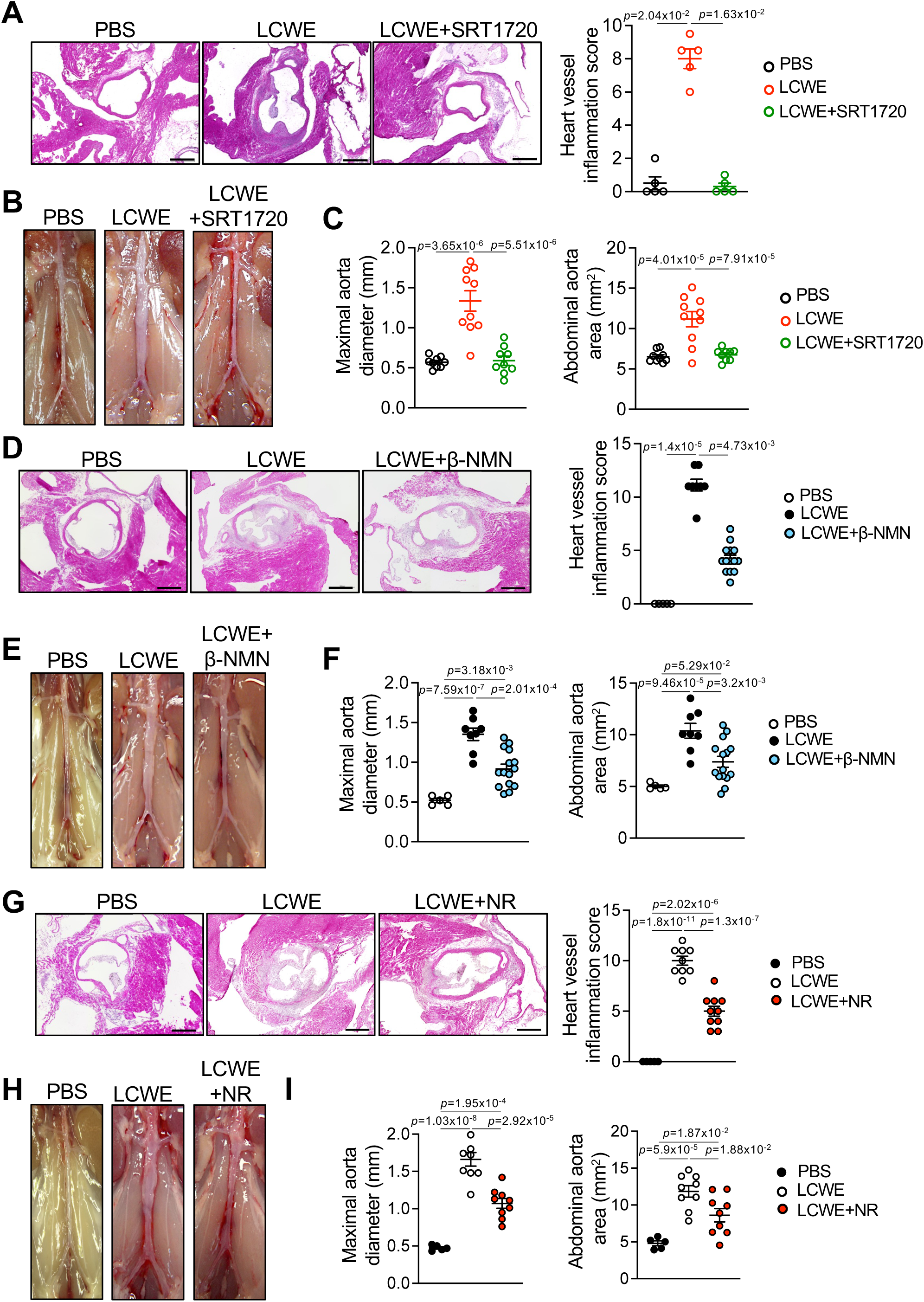
Boosting NAD^+^ levels or SIRT1 activity decreases *Lactobacillus casei* cell wall extract (LCWE)-induced KD severity. **(A-C)** WT mice were treated with SRT1720 i.p. (100mg/kg/body weight) daily, starting 1 day before until 7 days post LCWE injection and analyzed at 14 days. **(A)** Representative H&E-stained heart sections and heart vessel inflammation scores (n=5/group). Scale bars, 500μm. **(B)** Representative pictures of the abdominal aorta areas **(C)** Maximal abdominal aorta diameter and abdominal aorta area measurements (n=9 to 10/group). **(D-F)** WT mice were supplemented with β-NMN (500mg/Liter) in drinking water starting 1 day before LCWE injection and analyzed 7 days post-injection (n=8 to 15/group). **(D)** Representative H&E-stained heart sections and heart vessel inflammation scores. Scale bars, 500μm. **(E)** Representative pictures of the abdominal aorta areas **(F)** Maximal abdominal aorta diameter and abdominal aorta area measurements. **(G-I)** WT mice were treated with NR (200mg/kg/body weight) via oral gavage daily, starting 1 day before LCWE injection and analyzed 7 days post-injection (n=8 to 9/group). **(G)** Representative H&E-stained heart sections and heart vessel inflammation scores. Scale bars, 500μm. **(H)** Representative pictures of the abdominal aorta areas **(I)** Maximal abdominal aorta diameter and abdominal aorta area measurements. Data are presented as mean ± SEM. Pooled from 2 experiments (A-I). Each symbol represents one individual mouse. *\*p-value* < 0.05, ** *p-value* < 0.01, \*\*\**p-value* < 0.001, and \*\*\*\**p-value* < 0.001 by one-way ANOVA with Kruskal-Wallis test with Dunn’s multiple comparison tests (A, D), one-way ANOVA with Tukey’s post hoc test (C, F, G, I). β-NMN, β-Nicotinamide mononucleotide, NR, nicotinamide riboside.

## DISCUSSION

In this study, we demonstrate that LCWE-induced KD vasculitis impairs NAD^+^ metabolism and reduces the levels of the NAD^+^-dependent deacetylase SIRT1. Our data suggest that decreased NAD^+^ and SIRT1 levels contribute to the development of LCWE-induced cardiovascular lesions in mice. Furthermore, we establish the beneficial role of the NAD^+^-SIRT1 axis in this murine model of vasculitis, as deletion of *Sirt1* in VSMCs or myeloid cells exacerbates LCWE-induced cardiovascular inflammation, whereas pharmacological modulation of NAD^+^ metabolism or increasing SIRT1 expression and activity preventively or therapeutically is sufficient to decrease the severity of cardiovascular lesions in this murine model of KD.

The activation of Sirtuins mitigates the development of a broad array of cardiovascular diseases ^65^. SIRT1’s beneficial role in cellular regulation has been long established in the context of cardiovascular disease, particularly its modulation of mitochondrial function and reduction of oxidative stress ^66–68^. Consistent with previous research ^42,69,70^, our data demonstrate that SIRT1 activity enhances p62-LC3-II-dependent clearance of damaged mitochondria and inhibits mTOR-dependent suppression of autophagy during LCWE-induced KD, two pathways impaired in this model of vasculitis ^28^. Notably, patients with KD exhibit lower mRNA expression of autophagy-related genes, such as *LC3B*, *BECN1*, and *ATG16L1,* compared with febrile and HC ^71^. This impaired expression of autophagic genes recovers 21 days after IVIG treatment ^71^, highlighting the importance of intact autophagy in modulating the inflammatory response in KD. Indeed, supporting these findings, our analysis of publicly available datasets indicates decreased expression of several genes related to ROS metabolism and oxidative phosphorylation in patients with KD during the acute phase of the disease when compared with HC. We were able to detect a smaller number of genes related to autophagy/mitophagy, however, as those pathways are heavily modulated by posttranslational modifications and changes in protein abundance, mRNA levels might not accurately reflect protein levels during acute KD.

In addition to its role of promoting autophagy/mitophagy, SIRT1 has also been shown to deacetylate and activate PGC-1α (peroxisome proliferator-activated receptor gamma coactivator 1-alpha), the master regulator of mitochondrial biogenesis ^72^. Interestingly, we have not detected any changes in mtDNA levels, reflecting mitochondrial mass, in heart tissues of LCWE-injected *Sirt1^super^*mice. This could imply a balance between mitophagy and mitochondrial biogenesis during SIRT1 overexpression in LCWE-induced KD, as both pathways could potentially be upregulated to maintain a functional mitochondrial population during cardiovascular inflammation ^73.^

NAD^+^ can be used by SIRT1 as a co-substrate for its activity ^74^. Adults with low levels of SIRT1 are prone to early-onset microvascular dysfunction and may also face an increased risk of developing cardiovascular disease, which further highlights the importance of NAD^+^ metabolism in maintaining tissue homeostasis ^75^. Previous studies have shown that supplementation with NAD^+^ precursors reduces vascular oxidative stress and enhances mitochondrial and vascular functions ^55,76,77^. NAD^+^ precursors, NMN and NR, inhibit the production of IL-1β and TNF-α in a mouse model of angiotensin II-dependent endothelial inflammation ^78^. Boosting intracellular NAD^+^ levels promotes autophagy and rescues autophagic flux during ischemia/reperfusion (I/R) injury in rats, leading to decreased vascular damage, thereby preserving the heart microvasculature ^79^. Our data show that supplementing mice with NR or β-NMN decreases LCWE-induced KD severity, both preventively and therapeutically. Interestingly, while therapeutic supplementation with NR was effective in preventing both LCWE-induced heart inflammation and abdominal aorta dilation, NMN therapeutic supplementation reduced heart vessel inflammation, but not the abdominal aorta aneurysms. LCWE-induced abdominal aorta aneurysms develop acutely, starting around day 4 post-LCWE injection, and it is plausible that extracellular NMN needs to be dephosphorylated and converted to NR for cellular uptake and NAD^+^ synthesis ^80^, making NMN less effective therapeutically than NR when provided 3 days after LCWE injection. Interestingly, a GWAS on a cohort of children with KD from Korea identified a single-nucleotide polymorphism (SNP) in the *NMNAT2* gene as associated with increased susceptibility to KD ^81^. Although the downstream effects of this SNP have not been further studied, a dysfunctional *NMNAT2* gene may lead to defective NAD^+^ metabolism and potentially reduced SIRT1 activity in KD patients.

Longitudinal studies of the fecal microbiota from patients with KD indicate alterations in intestinal microbiota composition, associated with a decrease in relative abundances of bacteria known to produce short-chain fatty acids (SCFAs) ^82^. Among these bacteria, the abundance of *Akkermansia muciniphila* (*A. muciniphila*) is reduced in both KD patients and LCWE-injected mice that develop cardiovascular lesions ^82,83^. A connection between NAD^+^ metabolism and *A. muciniphila* was previously established in a murine model of amyotrophic lateral sclerosis (ALS), where supplementation of *A. muciniphila* ameliorated ALS symptoms in mice, and this beneficial effect was associated with the accumulation of nicotinamide in the central nervous system ^84^. Here, we show decreased circulating levels of NAD^+^-related metabolites in LCWE-injected mice, and we have previously demonstrated that oral supplementation with live *A. muciniphila*, which is decreased in LCWE-injected mice, reduces the severity of LCWE-induced KD vasculitis ^83^. While we have shown that the beneficial effect of *A. muciniphila* supplementation in LCWE-induced KD was mediated by SCFAs and Amuc_1100, a protein specific to the outer membrane of *A. muciniphila* ^83^, future studies will need to determine whether the accumulation of nicotinamide in cardiovascular tissues also contributes to the beneficial effect of *A. muciniphila* in LCWE-induced KD.

We observed enhanced mitochondrial coupling efficiency and acute response in mitochondria isolated from heart tissues of LCWE-injected *Sirt1*-overexpressing mice. These results could be attributed to SIRT1’s role in promoting mitochondrial biogenesis, expression of electron transport chain proteins ^85,86^, and the efficiency of oxidative phosphorylation coupled to ATP production ^72,87^. Indeed, in our proteomics analysis, we observed upregulation of multiple proteins involved in fatty acid oxidation and oxidative phosphorylation, suggesting that SIRT1 promotes substrate utilization by mitochondria to produce ATP. In addition, efficient mitochondrial respiration supports cardiovascular health by reducing oxidative stress ^43,88^. Under pathological conditions, there is excessive production of ROS, such as superoxide anion (O2^−^) and nitric oxide radical (NO). The increase in ROS surpasses the cellular antioxidant defenses, reducing the ability to scavenge ROS and contributing further to cardiac dysfunction and inflammation ^89,90^. Increased ROS production is associated with cardiovascular disease, such as atherosclerosis or I/R injury, where SIRT1 is protective by modulating antioxidant response ^91,92^. Consistent with these findings, we demonstrated that *Sirt1* overexpression is associated with reduced local ROS production in the hearts and abdominal aortas of mice, further mitigating cardiovascular inflammation in LCWE-induced KD.

Communication between stromal and immune cells is essential for a variety of biological functions, including immune responses to infections, tissue repair, and maintaining homeostasis, mediated mostly through cytokines and chemokines ^93^. Studies have reported that impaired autophagy in VSMCs contributes to abdominal aorta aneurysms ^94,95^. In a mouse model of dissecting aortic aneurysms induced by Angiotensin-II, VSMCs undergo clonal expansion and change into phagocytic-like cells, marked by increased expression of phagocytic, autophagic, and endoplasmic reticulum stress markers. In this model, inhibiting autophagy in VSMCs by deleting *Atg5* resulted in more severe and frequent aortic dissections, reduced autophagosome formation, and increased VSMC apoptosis ^96^. A published single-cell RNA sequencing analysis of the abdominal aortas from LCWE-injected mice reveals a phenotypic change in VSMCs, characterized by the emergence of proliferating type II VSMCs, which contribute to tissue inflammation and damage ^21^. This VSMCs phenotypic switch involved an increase in the expression of fibroblast-associated transcripts, MMPs, and transcripts associated with the recruitment of immune cells, concurrently with decreased expression of genes related to contractile functions ^21^. We now demonstrate that SIRT1 expression blocks the pathogenic type II switch of VSMCs in LCWE-induced cardiovascular lesions, as our proteomics analysis showed the upregulation of various VSMC contractile proteins with a downregulation of Mmp3 and Mmp14 in *Sirt1^super^* mice after LCWE injection, which was not observed in LCWE-injected WT control mice. This observation is consistent with the previous finding that *Sirt1* deletion in VSMCs in a mouse model of Marfan syndrome significantly increases MMP2 and MMP9 levels, which contribute to aortic wall remodeling and aneurysm formation and thereby aggravate thoracic aortic aneurysms ^97^. Furthermore, a reduction in SIRT1 protein levels was associated with vascular senescence and inflammation during angiotensin II-induced abdominal aortic aneurysms, and SIRT1 overexpression in VSMCs suppressed abdominal aortic aneurysm formation in mice ^98,99^. Deletion of *Sirt1* in VSMCs causes frequent elastin breaks and ROS production with severe aortic stiffness with angiotensin II treatment ^100^. Likewise, *Sirt1* overexpression or SRT1720 treatment counteracted oxidative stress during high-fat, high-sucrose diet-induced aortic stiffness ^101^. Consistent with these findings, our data indicate that specific deletion of *Sirt1* in VSMCs and macrophages exacerbates the severity of LCWE-induced cardiovascular lesions and boosts the production of IL-1β and TNFα by macrophages.

In LCWE-induced KD, myeloid cell-derived IL-1β contributes to the development of cardiovascular lesions ^16,22^. Here, we show that *Sirt1* overexpression reduced IL-1β levels *in vivo* and *in vitro*. Furthermore, macrophage-specific *Sirt1* deletion increased the severity of LCWE-induced cardiovascular lesions and was associated with enhanced *in vivo* IL-1β production. Similarly, BMDMs isolated from the same mice secreted more IL-1β and TNF-α upon LCWE stimulation *in vitro*. Additionally, caspase-1 activity was higher in cardiovascular tissues of mice lacking SIRT1 in their myeloid cells after LCWE injection, hinting at higher NLRP3 activity and production of IL-1β. Therefore, *Sirt1* deletion enhances the production of proinflammatory cytokines from macrophages, contributing to the development of cardiovascular lesions. Consistently, BMDMs lacking *Sirt1* were more prone to proinflammatory M1-like polarization, further confirming that *Sirt1* activity counteracts inflammatory cytokine production from macrophages. These findings were consistent with previous studies showing that SIRT1 inhibits inflammatory M1 polarization and promotes M2 phenotype of macrophages during arthritis and acute lung injury ^102,103^.

KD is the leading cause of acquired heart disease among children ^6^. Since up to 20% of patients with KD do not respond to IVIG treatment and face a higher risk of cardiovascular lesions during childhood and long-term consequences in adulthood ^104^, discovering new therapeutic interventions is crucial. Altogether, our data identify SIRT1 as a beneficial modulator during LCWE-induced KD. Mechanistically, SIRT1 promotes clearance of damaged mitochondria in VSMCs and prevents their switch to a more proliferative and pathogenic phenotype, and reduces proinflammatory cytokine production from macrophages and their polarization towards an M1-type inflammatory state. Our results indicate that boosting NAD^+^ metabolism and downstream SIRT1 activity can be a potential new therapeutic strategy to prevent or treat cardiovascular inflammation linked to KD vasculitis. Targeting autophagy via the NAD^+^-SIRT1 axis in VSMCs could be highly effective, since there are no established treatments that directly target the development of luminal myofibroblast proliferation or long-term vascular complications in KD patients. Nonetheless, further studies are necessary to confirm the applicability of our findings in patients with KD.

## MATERIALS and METHODS

### Mice

All procedures were approved by the Institutional Animal Care and Use Committee of Cedars-Sinai Medical Center. Mice were housed at 22 °C with a 12 h light/12 h dark cycle and fed with a regular chow diet. Wild-type (WT) C57BL/6J, 129S1/SvImJnd (#002448), *Sirt1^super^* (#024510), *Sirt1^ER/ER^* (#043520), *Myh11^Cre^*^/ERT^ (#019079), and *Csfr1^cre^* (#029206) mice were obtained from the Jackson Laboratory (Bar Harbor, ME, USA). *Sirt1^fl/fl^* (Jackson Laboratory #029603) were a kind gift from Dr. Changfu Yao at Cedars-Sinai Health Sciences Institute and were crossed with *Myh11^Cre^*^/ERT2^ mice and *Csfr1^cre^* mice in our colony; littermates were used as controls. Mice were housed under specific pathogen-free conditions at Cedars-Sinai Medical Center and provided with a standard diet and water *ad libitum* unless noted otherwise for experimental purposes. Five week-old male mice were used for experimental purposes, as LCWE-injection induces stronger and more consistent coronary vasculitis lesions and abdominal aorta aneurysms in male mice than female mice ^6,23^. Mice were randomly assigned to the different experimental groups.

### The Lactobacillus casei cell wall extract (LCWE)-induced KD vasculitis murine model

*Lactobacillus casei* (ATCC; 11578) cell wall extract was prepared as previously described ^16^. Five week-old male mice were injected i.p. with 500mg of LCWE or PBS (control group). One or 2-weeks post-injection, mice were euthanized, and blood was collected. Hearts and aortas were harvested and embedded in Optimal Cutting Temperature (OCT, Sakura Finetek) compound for histology. Abdominal aortas were photographed after dissection and then embedded in OCT. Maximal abdominal aorta diameter was calculated by measuring 5 different areas on the infrarenal section with ImageJ (NIH). The infrarenal abdominal aorta area was also measured in ImageJ. Representative images were chosen based on corresponding inflammatory scores or aorta measurements reflecting the mean of the sample group.

### Tissue fixation, H&E staining, and histological analysis

7μm-thick serial cryosections from OCT-embedded heart tissues were stained with hematoxylin and eosin (H&E, Sigma-Aldrich) for histological analysis or were used for immunofluorescent staining. Histopathological examination and inflammatory scoring of heart tissues were performed from these sections by a senior investigator blinded to the experimental groups. Heart vessel inflammatory score (coronary arteritis, aortic root vasculitis, and myocarditis) was determined as previously described ^16^. Only heart sections showing the coronary artery (CA) branch separating from the aorta were scored. Briefly, acute inflammation, chronic inflammation, and connective tissue proliferation were each assessed using the following scoring system: 0 = no inflammation, 1 = rare inflammatory cells, 2 = scattered inflammatory cells, 3 = diffuse infiltrate of inflammatory cells, and 4 = dense clusters of inflammatory cells. Fibrosis was determined using the following scoring system: 0 = no medial fibrosis, 1 = medial fibrosis involving less than 10% of the CA circumference, 2 = medial fibrosis involving 11% to 50% of the CA circumference, 3 = medial fibrosis involving 51% to 75% of the CA circumference, and 4 = medial fibrosis involving more than 75% of the CA circumference. All 4 scores were combined to generate a severity score called “Heart vessel inflammation score”, as previously published ^16^. Pictures were taken using Biorevo BZ-9000 or BZ-X710 microscope (Keyence). Representative images were chosen based on corresponding inflammatory scores that were closest to the mean of the sample group.

### Inducible deletion of Sirt1 in smooth muscle cells (SMCs)

To delete *Sirt1* in SMCs, *Myh11^Cre^*^/ERT2^ *Sirt1^Δ/Δ^* and *Sirt1^fl/fl^* littermate controls were fed with a tamoxifen diet (250mg/kg, Envigo; TD.130856) for 2 weeks, then regular chow for 4 weeks prior to LCWE injection. Two weeks after LCWE injection, hearts and abdominal aortas were harvested, and the severity of LCWE-induced KD vasculitis was determined as above.

### Inducible induction of Sirt1-ER expression in mice

To induce the expression of SIRT1-ER protein, *Sirt1^ER/ER^*mice and 129S1/SvImJnd WT controls were injected i.p. 75 mg/kg tamoxifen for 5 consecutive days starting 1 week after LCWE injection. At the end of 5 days, mice were rested for another week until the tissues were harvested. To quantify IL-1β in the peritoneal lavage, WT and *Sirt1^ER/ER^* mice were first injected i.p. with 75 mg/kg tamoxifen for 5 consecutive days, then injected with LCWE. 24 hours later, peritoneal lavage was collected in sterile PBS.

### IL-1β quantification from mice peritoneal lavage

Mice were injected with either PBS or LCWE as described above. 24 hours later, peritoneal lavage was collected in sterile PBS, centrifuged at 400*g* at 4°C for 5 minutes, and the cells were discarded. IL-1β concentration was assessed from the supernatants by ELISA (R&D Systems, Mouse IL-1 beta/IL-1F2 DuoSet ELISA; DY401) according to the manufacturer’s protocol.

### In vivo SRT1720, Resveratrol, NR, and β-NMN treatments

WT mice were either injected with SRT1720 (100 mg/kg, i.p., InvivoChem; V0428), or orally treated with Resveratrol (200 mg/kg in 0.5% w/v of methyl cellulose and 0.2% Tween80, homogenized and filtered, daily oral gavage, Sigma-Aldrich; R5010), NR (200mg/kg, daily oral gavage, Fisher Scientific; NC1611436) or β-NMN (500mg/L in drinking water, Santa Cruz Biotechnology; sc-212376B). Mice were euthanized at the experimental endpoint, and tissues were harvested and analyzed as described above. In another set of experiments, WT mice were first injected with LCWE and then supplemented with either β-NMN or NR on day 3 post-LCWE injection, as described above.

### In vivo autophagic flux assay

WT mice were first injected with LCWE, and 2 weeks later were injected i.p. with a single dose of chloroquine (CQ) at 40mg/kg body weight. Four hours after CQ injection, abdominal aortas were collected, and total cell lysates were subjected to Western blot analysis.

### NAD^+^ measurement

NAD^+^ levels from serum were quantified using the NAD/NADH assay kit (Abcam; ab65348) according to the manufacturer’s protocol and reported as relative levels to control samples.

### Primary smooth muscle cell isolation and culture

Mouse primary coronary artery smooth muscle cells (Cell Biologics) were grown in DMEM supplemented with 10% fetal bovine serum (FBS) 1% L-glutamine, and 1% penicillin/streptomycin cocktail (P/S) in a humidified, 5% CO2 incubator at 37 °C. Where indicated, cells were treated with recombinant IL-1β (rIL-1β; 10ng/ml; R&D Systems; 401-ML-010/CF), SRT1720 (5μM; InvivoChem; V0428), EX527 (5μM; InvivoChem; V0429), Chloroquine (5μM; CQ, Sigma-Aldrich; C6628) and LCWE (60 μg/ml) for 24 hours. To isolate VSMCs, abdominal aortas from *Sirt1^fl/fl^* and *Myh11^Cre/ERT2^Sirt1^Δ/Δ^* mice were perfused with Hank’s Balanced Salt Solution (HBSS, Fisher Scientific Gibco™ HBSS, calcium, magnesium, no phenol red) and harvested below the left renal artery to the iliac arteries. 2-3 aortas per group were pooled and incubated for 10 minutes in enzyme solution 1 at 37 °C and 5% CO_2_ (1 mg/ml collagenase type II (Worthington Biochemical; LS004174), 0.744 units/ml elastase (Worthington Biochemical; LS0022792), and 1% P/S). Adventitia was stripped off under the microscope and aortas were further incubated with enzyme solution 2 at 37 °C and 5% CO_2_ for 1 hour (1 mg/ml collagenase type II, 2 units/ml elastase and 1% P/S). After an hour, cells were triturated with a Pasteur pipette, washed twice with DMEM/F12 media, and centrifuged at 300 *g* for 5 minutes. Cells were plated into 3 wells of a 48 well-plate in DMEM/F12 with 20% FBS and 1% P/S and left undisturbed for a week. Where indicated, cells were stimulated with 10ng/ml rIL-1β for 24 hours. To induce SIRT1 deletion, primary smooth muscle cells isolated from *Sirt1^fl/fl^* and *Myh11^Cre/ERT2^Sirt1^Δ/Δ^* mice were treated with 1μM 4-Hydroxytamoxifen (4-OHT; MedChemExpress HY-16950) for 48 hours.

### Bone marrow-derived macrophages (BMDMs)

Bone marrows were collected from bilateral tibia and femurs of mice with RPMI containing 1% P/S cocktail and filtered through a cell strainer (BD; 352350). Cells were centrifuged at 300 g for 5 min and resuspended in RPMI enriched with 20% L929 cells conditioned medium, 10% FBS, and 1% P/S cocktail, followed by growth on Petri dishes and differentiation to macrophages for 7 days. After differentiation, cells were treated with LCWE (60 μg/ml) or LPS (10ng/ml; Invivogen, tlrl-3pelps) together with IFN*γ* (50ng/ml; R&D Systems, 485-MI-100/CF) for M1 polarization or IL-4 (10ng/ml; Tonbo Biosciences, 21-8041-U020) for M2 polarization for 24 hours. For cytokine measurements, supernatants were collected to measure IL-1β (R&D Systems, Mouse IL-1 beta/IL-1F2 DuoSet ELISA; DY401) and TNF-α levels by ELISA (eBioscience; 50-137-02), according to the manufacturer’s instructions.

### RNA isolation and quantitative real-time PCR analysis (qRT-PCR)

Total RNA was isolated by TRIzol Reagent (Invitrogen; 15596018) according to manufacturer’s protocol and reverse transcribed by using RevertAid First strand cDNA synthesis kit (ThermoScientific; K1691) to complementary DNA (cDNA), according to manufacturer’s protocol. cDNAs were amplified using specific primers specified below and Power-Up-SYBR green (Applied Biosystems; A25742) on a CFX96 Real-Time System (BioRad). Gene expression was quantified using triplicate samples with the relative threshold ΔΔCt method: ΔΔCt = (primer efficiency)^(−ΔΔCt), where ΔΔCt means ΔCt (target gene) −ΔCt (reference gene).

### Protein lysates, SDS/PAGE electrophoresis, transfer, and Western blotting

Cells or tissues were lysed in RIPA buffer (0.5M Tris-HCl, pH 7.4, 1.5M NaCl, 2.5% deoxycholic acid, 10% NP-40, 10mM EDTA, Millipore; 20-188). Lysates were cleared with brief centrifugation for 10 min at 8000*g*, normalized, and boiled at 95 °C after addition of 6X SDS loading dye. Proteins were then loaded to gradient SDS/PAGE gels and transferred to nitrocellulose or PVDF membranes. Blocking (1 hour at room temperature) and primary (overnight at 4°C), followed by secondary antibody (1 hour at room temperature) incubation in 5% (w/v) dry milk or bovine serum albumin (in tris-buffered saline buffer with 0.1% Tween-20 (v/v)) on a shaker. Membranes were developed in ECL prime reagent (ThermoFisher Scientific; 34577) and images were captured with ChemiDoc Imager (BioRad). Blots shown are representative of three or more experiments. Quantifications were performed with Image Lab software (BioRad).

### Mitochondrial enrichment

Cells were homogenized in mitochondrial isolation buffer (250 mM sucrose, 1 mM EDTA, 10 mM HEPES, pH 7.4) with inhibitors (1X phosphatase inhibitor cocktail 3 and 1X protease inhibitor cocktail, (Sigma-Aldrich; P0044 and P8340) by running through 27.5 g needle three times. Nuclei and cell debris were eliminated by low-speed spin down (600*g*, 4 °C, 5 min). A portion of the supernatant was saved as cell lysate, and the rest was further centrifuged (7000*g*, 4 °C, 15 min) to obtain the final mitochondria-enriched pellet and supernatant (cytosol). The mitochondria-enriched pellet was resuspended in isolation buffer and centrifuged (600*g*, 4 °C, 5 min) as a final wash. The pellet was resuspended in cold RIPA buffer with inhibitors. Protein concentrations were measured with DC Protein Assay Kit II (BioRad; 500-0112). Normalized samples are boiled in SDS loading dye at 95 °C for 5 min before loading on gradient SDS/PAGE gels. Samples were then transferred to nitrocellulose membranes and stained with Ponceau S solution (Sigma-Aldrich; P7170-1L), washed with distilled water, and imaged as loading control before the blocking step.

### Assessment of mitochondrial DNA (mtDNA) copy number

Total DNA from heart tissues was isolated using *Quick*-DNA Miniprep Plus Kit (Zymo Research; D4068) according to manufacturer’s protocol. DNA (0.2 μg) was amplified using nuclear- or mitochondria-encoded genes-specific primers using Power-Up-SYBR green (Applied Biosystems; A25742) on a CFX96 Real-Time System (BioRad). mtDNA: nuclear DNA ratios were calculated by normalizing results of mitochondria-encoded (*Cox1* or *Nd4*) gene to nuclear-encoded (*ApoB*) genes.

### Oxygen consumption rate (OCR) and extracellular acidification rate (ECAR) measurements

Freshly isolated mitochondria from heart tissues ^105^, were used to perform mitochondrial respirometry assay using Seahorse XF Cell Mito Stress Test Kit (Agilent; 103015-100) according to the manufacturer’s instructions. XFe^96^ well plates were loaded in port A with ADP (0.25 mM final concentration), pyruvate (10 mM final concentration), and malate (2 mM final concentration), port B with oligomycin (2 µM final concentration), port C with carbonyl cyanide 4-(trifluoromethoxy) phenylhydrazone (FCCP, 2 µM final concentration), and port D with antimycin A/rotenone (1 µM final concentration for each). Measurements were acquired with Seahorse XFe^96^ Analyzer (Agilent Technologies) and analyzed with WAVE software (Agilent Technologies).

### Immunofluorescent staining

7μm thick sections were obtained from OCT frozen heart and abdominal aortas, fixed in ice-cold acetone for 5 minutes, washed with PBS and blocked 1 hour at room temperature with 3% donkey serum (Abcam; ab7475) in PBS. Sections were then incubated with primary antibodies (SIRT1, p62, LC3B, Lumican, Alexa Fluor 488 anti-alpha-smooth muscle actin, Alexa Fluor 647 anti-mouse F4/80, NLRP3) or isotype controls in blocking solution overnight at 4°C in a humid chamber. The next day, sections were washed again with PBS, incubated 1 hour at room temperature with fluorescently labeled secondary antibodies (anti-mouse or rabbit Alexa Fluor 594 or Alexa Fluor 488) prepared in blocking solution, followed by another washing step with PBS. Slides were mounted with fluoroshield mounting medium with DAPI (Abcam; ab104139). Caspase-1 activity in tissues was detected using the fluorochrome-labeled inhibitors of caspases assay (FLICA) (Immunochemistry Technologies; Cat #SKU 98), according to the manufacturer’s protocol. Images were obtained using a Biorevo BZ-X710 (Keyence) fluorescent microscope and were further analyzed with ImageJ (NIH) software. For all immunofluorescent stainings, isotype controls for all primary antibodies were used to distinguish positive staining from the background.

### MitoTracker and mitoSOX staining

MitoTracker Green FM (Invitrogen; M7514) and mitoSOX red (Invitrogen; M36008) were used to stain mitochondria and mitochondrial ROS, respectively, according to the manufacturers’ protocols. After staining, cells were fixed with 4% paraformaldehyde, mounted onto slides with Fluoroshield mounting medium with DAPI (Abcam; ab104139), visualized with Biorevo BZ-X710 (Keyence) fluorescent microscope and mean fluorescence intensity was calculated in ImageJ software.

### Dihydroethidium (DHE) staining for ROS measurement

Frozen sections of heart tissues were stained with DHE (Sigma-Aldrich; D7008) to quantify ROS production, as previously published ^106^. Integrated density of the fluorescent signal was calculated from five randomly selected equal sized areas in each tissue section with ImageJ (NIH).

### Proteomics analysis of abdominal aorta tissues

#### Tissue preparation

Tissue samples were prepared in parallel using the S-Trap 96-well plate (Protifi, C02-96-well-10) following the manufacturer’s protocol. Briefly, the tissue was homogenized using a probe polytron, and the proteins were reduced using 50 mM dithiothreol for 30 min at 37°C and subsequently alkylated with 20 mM iodoacetamide (IAA) for 30 min at RT in the dark, before acidification with phosphoric acid (1.2% final concentration). The samples were diluted 1:7 with binding/wash buffer (90% methanol, 100 mM TEAB, pH 7.4), loaded on the S-Trap 96-well plate and centrifuged for 5 min at 1500*g* (RT). The columns were washed three times with 200 µL binding/wash buffer and centrifuged each time for 5 min at 1500*g* (RT). One microgram of trypsin per 125 µL 50 mM TEAB (pH 7.4) was added to the columns and digestion was performed overnight (approximately 16 h) at 37°C. Peptides were eluted in three consecutive centrifugation steps (1500*g* for 5 min at RT) using 80 µL 50 mM TEAB (pH 7.4), 80 µL 0.2% formic acid in dH2O and 80 µL 2% ACN, 0.1% formic acid in dH_2_O. The eluates were vacuum dried and resuspended in 0.1% formic acid in dH_2_O for MS analysis.

#### LC-MS/MS analysis

DIA analysis was performed on an Orbitrap Exploris 480 (Thermo Scientific) mass spectrometer interfaced with a microflow-nanospray electrospray ionization source (Newomics, IS-T01) coupled to Vanquish Neo ultra-high-pressure chromatography system with 0.1% formic acid in water as mobile phase A and 0.1% formic acid in acetonitrile as mobile phase B. Peptides were separated at an initial flow rate of 15 µL/minute and a linear gradient of 4-28% B for 0-52 minutes, 28-40% B for 52-55 minutes. The column was then flushed with 98% B for 55-60 minutes before being re-equilibrated for the next run. The column used was Thermo Scientific™ µPac™ HPLC column with a 200cm bed length (P/N: COL-NANO200G1B). Source parameters were set to a voltage of 3500 V and a capillary temperature of 320°C. MS1 scan range was set to 300-1200 m/z and MS1 resolution was set to 120,000 with an AGC target set to ‘Standard’. RF Lens was set to 50% with maximum injection time set to ‘Auto’. Fragmented ions were detected across a scan range of 400-1000 m/z with 50 non-overlapping data-independent acquisition precursor windows of size 12 Da. MS2 resolution was set to 15,000 with a scan range of 200-2000 m/z and normalized HCD collision energy set to 32%. Maximum injection time mode was set to ‘Custom’ with a maximum injection time of 24 ms. All data is acquired in profile mode using positive polarity.

#### Data analysis

MS raw data files were searched against UniProt mouse reviewed protein sequence entries (accessed April 2023) using DIA-NN (v 1.8.1)^107^ in library free mode with default parameters. Based on recent comparisons with library-based approaches, DIA-NN in library-free mode has been found to produce results that are comparable or better than those of experimental library-based searches while being freely available and was hence chosen for the analysis of all data^108^. The output protein group matrix from DIA-NN was used to perform downstream analysis using MetaboAnalyst 6.0^109^.

#### Bioinformatics

Filtration of the 5777-signal DIA-NN-normalized protein matrix was performed prior to downstream analysis. Intensity signals corresponding to multiple or duplicate proteins were omitted, removing 802 signals from the data. Proteins without an intensity signal in at least 1 replicate for each condition were omitted from downstream analysis as well, removing 830 signals, leaving a filtered 4145-signal DIA-NN protein matrix. Missing values were imputed with 20% of the minimum intensity signal for the given protein, and the data was then log2-transformed. The data was uploaded to MetaboAnalyst for differential expression analysis. Three comparisons were tested: WT LCWE vs. WT PBS, SIRT1 TG LCWE vs. SIRT1 TG PBS, and SIRT1 TG LCWE vs. WT LCWE. The 443 overlapping differentially expressed proteins between WT LCWE vs. WT PBS and SIRT1 TG LCWE vs. WT LCWE were identified and analyzed for significant functional networks with ClueGO^110^ utilizing Gene Ontology’s Biological Process database, separating proteins by upregulation and downregulation in WT LCWE vs. WT PBS (Statistical Test Used: Enrichment/Depletion, Use *p-value* cutoff: True, *p-value* cutoff: 0.05, Correction Method Used: Bonferroni step down, Min GO Level: 3, Max GO Level: 5, Number of Genes: 10, Get All Percentage: true, GO Fusion: true, GO Group: true, Kappa Score Threshold: 0.4, Over View Term: SmallestPValue, Group By Kappa Statistics: true, Initial Group Size: 1, Sharing Group Percentage: 50.0). Pathway analysis on the overlapping genes was done with Qiagen Ingenuity Pathway Analysis^111^ for identification of pathways significantly activated (Activation z-score>2 & *adj. p-value*<0.05) or inhibited (Activation z-score<-2 & *adj. p-value*<0.05) in the 3 comparisons. Heatmaps were generated using the heatmap R package, and PCA, bubble and bar plots with created with the ggplot2 R package.

### Analysis of human gene expression data sets

Publicly available gene expression data sets GSE68004^112^ and GSE73461^113^ were obtained from National Center for Biotechnology Information’s Gene Expression Omnibus (GEO; https://www.ncbi.nlm.nih.gov/geo/). Transcriptomic datasets were accessed and loaded into R via GEOQuery, and summary statistics were generated with limma topTable function. Experimentally observed genes involved in Oxidative Phosphorylation and Autophagy/ROS response were queried from Gene Ontology’s database AmiGO 2. 119 genes involved in oxidative phosphorylation and 448 genes involved in autophagy/ROS metabolism were retrieved. The expression of these genes was assessed in whole blood of patients with acute KD and HCs (n = 37 HCs and n = 76 acute KD patients for GSE68004^112^; and n = 55 HCs and n = 77 acute KD patients for GSE73461^113^). Differentially expressed genes (DEGs) (Benjamini-Hochberg–*adj. p-value*< 0.05 and FC > 1.5) from the gene signatures were identified in these 2 data sets (GSE68004 and GSE73461).

### Metabolomics and data analysis

The metabolomic analysis was performed by Metabolon, Inc (Durham, NC, USA). Serum samples were collected from WT mice at baseline (Day 0) and at 2 weeks post-LCWE injection. Samples were prepared using the Metabolon proprietary automated system, and obtained extracts were analyzed using an ultra-high performance liquid chromatography-tandem mass spectrometry (UPLC-MS/MS) platform. Metabolites were identified by matching their chromatographic retention time and mass spectra within Metabolon’s proprietary library. Metabolite names and scaled peak intensity data were provided by Metabolon^114^, and visualized using R and Metaboanalyst. Welch’s two-sample t-test was used to identify Metabolites that differed significantly between experimental groups.

### Statistical analysis

GraphPad Prism 10 (GraphPad Software, Inc) or R software was used to statistically analyze data and create graphs. Statistical analyses used for each figure panel are indicated in the figure legends. For comparisons of 2 groups, a 2-tailed unpaired Student’s t-test, with Welch’s correction when indicated, was used for normally distributed data. For nonparametric data, the Mann-Whitney two-tailed U/Wilcoxon rank test was used. For more than 2 group comparisons, one-way ANOVA with Tukey post-test analysis was used for normally distributed data. The Kruskal-Wallis test with Dunn’s multiple comparisons test was used for non-normally distributed data. A two-way ANOVA was used when there were two independent variables with multiple groups, such as genotype and treatment. Results are reported as means ± SEM, where each point represents one sample. A *p-value* <0.05 was considered statistically significant. No statistical methods were used to predetermine the sample size.

## ACKNOWLEDGEMENTS

We thank Dr. Changfu Yao for kindly gifting us the *Sirt1^fl/fl^* mice. Graphical abstract and experimental schematics were created with Biorender.com.

## SOURCES OF FUNDING

A.E. Atici is supported by the American Heart Association Postdoctoral Fellowship 1196203. Work in this study was supported by the National Institutes of Health (NIH) grant R01HL159297 to M. Noval Rivas. M. Noval Rivas is supported by the R01 HL139766 and R01 HL159297. M. Arditi is supported by the NIH grants R01 HL170580 and NIH R01 HL149972. M. Arditi and M. Noval Rivas are supported by the NIH R01AI157274 grant.

## AUTHOR CONTRIBUTIONS

A.E. Atici and M. Noval Rivas conceived the project and designed the experiments. A.E. Atici, P.K. Jena, T.T. Carvalho, B. Ross, E. Aubuchon, M.E. Lane, A.C. Gomez, Y. Lee performed experiments. M. Arditi, S. Chen and T.R. Crother helped conceptualize and discuss results. A.E. Atici and M. Noval Rivas wrote the article with input from all authors.

## DISCLOSURES

The authors report no conflict of interest.

## NOVELTY AND SIGNIFICANCE

### What is known?

- Sirtuin 1 (SIRT1) is an NAD^+^ dependent deacetylase with anti-inflammatory function, which promotes autophagy/mitophagy pathways in the context of cardiovascular diseases.
- Kawasaki disease (KD) is an acute systemic pediatric vasculitis and is the leading cause of acquired heart disease. Expression of autophagy-related genes has been shown to decrease in KD patients during the acute phase and is recovered after intravenous immunoglobulin G (IVIG) treatment.
- The *Lactobacillus casei* cell wall extract (LCWE)-induced murine model of KD vasculitis replicates key characteristics of the human disease, and autophagy/mitophagy pathways are impaired in cardiovascular tissues of LCWE-injected mice.

### What New Information Does This Article Contribute?

- We uncover a dysfunctional SIRT1-NAD^+^ axis as well as impaired downstream SIRT1 signaling in a murine model of KD.
- Boosting SIRT1 activity in smooth muscle cells induces autophagy/mitophagy, promotes the clearance of damaged mitochondria, reduces reactive oxygen species (ROS) formation, increases mitochondrial respiration, and counteracts pro-inflammatory cytokine production of macrophages during cardiovascular inflammation linked to LCWE-induced KD.
- Boosting NAD^+^ metabolism by supplementing mice with NAD^+^ precursors or pharmacological activation of SIRT1 *in vivo* emerges as a novel therapeutic target for KD.

Intact autophagy/mitophagy is critical for homeostasis in tissues that are highly dependent on mitochondrial function to produce energy, such as the cardiovascular system. Impaired removal of damaged or dysfunctional mitochondria contributes to mitochondrial ROS formation, jeopardizes ATP-linked oxidative phosphorylation, and exacerbates inflammation during cardiovascular disease. KD is the leading cause of acquired heart disease in children and causes coronary artery aneurysms if untreated. Dysfunctional autophagy/mitophagy has been reported in patients with KD; however, the mechanism behind this immunopathology is unknown. Here, we observed that NAD^+^ production and SIRT1 levels are decreased in a murine model that mimics the main features of KD. Overexpression of SIRT1 in mice decreased the severity of vasculitis, where its deletion in smooth muscle or myeloid cells promoted cardiovascular inflammation and vasculitis in this murine model of KD. Mechanistically, SIRT1 overexpression counteracts impaired autophagy/mitophagy pathways, reduces ROS production, and enhances mitochondrial respiration *in vivo*. Furthermore, boosting NAD^+^ levels with precursors therapeutically or preventatively and activating SIRT1 pharmacologically decreased cardiovascular lesions in this murine model of KD. These findings highlight a dysfunctional NAD^+^-SIRT1 axis in KD and identify modulation of NAD^+^ levels or SIRT1 activity as novel therapeutic approaches to improve the related cardiovascular outcomes.

## NONSTANDARD ABBREVIATIONS AND ACRONYMS

BMDM: bone marrow-derived macrophage
CAA: coronary artery aneurysm
CQ: chloroquine
DHE: dihydroethidium
ECAR: extracellular acidification rate
FLICA: fluorochrome-labeled inhibitors of caspases
IL: interleukin
IVIG: intravenous immunoglobulin G
KD: Kawasaki disease
LCWE: *Lactobacillus casei* cell wall extract
NMN: nicotinamide mononucleotide
mtDNA: mitochondrial DNA
NR: nicotinamide riboside
OCR: oxygen consumption rat
RES: resveratrol
ROS: reactive oxygen species
WT: wild-type

## SUPPLEMENTARY FIGURE LEGENDS

**Figure S1. Metabolomic analysis revealed a deficit in the NAD^+^ pathway during LCWE-induced KD. (A)** Schematic of the experimental design. Blood was collected from WT mice at day 0 (baseline) and 14 days after LCWE injection for metabolomics analysis. (**B**) Schematic of the different pathways involved in NAD^+^ production, depletion, and consumption. NAD^+^, nicotinamide adenine dinucleotide, NR, nicotinamide riboside, NMN, nicotinamide mononucleotide, NAM, nicotinamide, NRK, nicotinamide riboside kinase, NAMPT, nicotinamide phosphoribosyltransferase, NAMNAT, nicotinamide/nicotinate mononucleotide adenylyltransferase, QA, quinolinic acid, QPRT, quinolinate phosphoribosyl transferase, NA, nicotinic acid, NAMN, nicotinic acid mononucleotide, NAPRT, NA phosphoribosyltransferase, NMNAT, NMN adenylyltransferases, NADS, NAD synthase, PARPs, poly (ADP-ribose) polymerases.

**Figure S2. SIRT1 protein levels in heart tissues of transgenic mice. (A)** Protein lysates of the apex part of heart tissues from WT and *Sirt1^super^* mice injected with LCWE were analyzed by Western blotting using specific antibodies for SIRT1 and β-actin (n=5/group). Representative of 2 experiments. **(B)** Levels of IL-1β in the peritoneal lavage of WT and *Sirt1^ER/ER^* mice injected 5 consecutive days of tamoxifen (75mg/kg) and then with PBS or LCWE 24 hours post-injection (n=5/group). \**p-value*<0.05, \*\**p-value*<0.01, \*\*\**p-value*<0.001, and \*\*\*\**p-value*<0.001 by two-way ANOVA with Tukey post hoc test (B).

**Figure S3. SIRT1 overexpression counteracts LCWE-induced KD related inflammation in abdominal aortas. (A)** Schematic of the experimental design. Abdominal aortas of either PBS or LCWE-injected WT and *Sirt1^super^* mice were collected 2 weeks post-injection (n=5/group), and tissue processed for untargeted proteomics analysis. **(B)** Heatmap of global protein expression data from abdominal aortas of either PBS or LCWE-injected WT and *Sirt1^super^* mice 2 weeks post-injection (n=5/group). **(C)** ClueGO Ontology analysis of pathways from LCWE-induced KD DEPs the highlighted proteins in red from (B), which are significantly increased in LCWE-injected WT mice compared with PBS-injected WT mice and downregulated in LCWE-injected *Sirt1^super^* mice compared with LCWE-injected WT mice (n=5/group). **(D)** ClueGO Ontology analysis of pathways from the highlighted proteins in red from (B), which are significantly decreased in LCWE-injected WT mice compared with PBS-injected WT mice and increased in LCWE-injected *Sirt1^super^* mice compared with LCWE-injected WT mice (n=5/group).

**Figure S4. SIRT1 increases mitochondrial respiration efficiency but does not impact mtDNA levels during LCWE-induced KD (A)** Oxygen consumption rate (OCR) in isolated mitochondria of heart tissues from LCWE-injected WT or *Sirt1*^super^ mice at 2 weeks post-injection. (n=5/group). **(B)** Extracellular acidification rate (ECAR) in isolated mitochondria of heart tissues from LCWE-injected WT or *Sirt1*^super^ mice at 2 weeks post-injection. (n=5/group). Oligomycin: ATP synthase inhibitor; FCCP: mitochondrial uncoupler; R/A: rotenone and antimycin A mix (inhibitors for ETC complex I and III, respectively). **(C)** qRT-PCR quantification of total genomic DNA and mtDNA (*Cox1* and *Nd4*) to nucDNA (*ApoB*) ratio from heart tissues of WT and *Sirt1^super^* mice injected with LCWE 2 weeks post-injection (n=5/group). Data are presented as mean ± SEM. Each symbol represents one individual mouse. \**p-value*<0.05, \*\**p-value*<0.01, \*\*\**p-value*<0.001, and \*\*\*\**p-value*<0.001 by unpaired t-test (C). Pyr/Mal, pyruvate/malate, ADP, adenosine diphosphate, FCCP, Carbonyl cyanide p-trifluoromethoxyphenylhydrazone.

**Figure S5. Expression of signature genes involved ROS metabolism and oxidative phosphorylation in patients with KD. (A-B)** Heatmaps illustrating the expression of genes involved in ROS metabolism (A) and oxidative phosphorylation (B) in whole blood from patients with KD compared with healthy control (HC) (GSE68004 and GSE73461).

**Figure S6. Overexpression of SIRT1 reduces ROS accumulation and promotes autophagic flux during LCWE-induced KD. (A)** Representative dihydroethidium (DHE) staining heart (upper) and abdominal aorta (lower) tissue cross-sections collected from WT and *Sirt1^super^* mice injected with PBS or LCWE at 2 weeks post-injection (n*=*5/group). Scale bars: 50 µm. **(B)** Western blot quantification of SIRT1 and pUbiquitin^S65^ in protein lysates of heart tissues from WT and *Sirt1^super^* mice injected with LCWE at 2 weeks post-injection (n=4, 5/group). **(C)** Schematic of the *in vivo* autophagic flux assay experimental design. WT and *Sirt1^super^*mice were injected with LCWE, and 14 days later received chloroquine (CQ) i.p. 4 hours before collecting the abdominal aortas. **(D)** Representative Western blots of p62, LC3-I and LC3-II protein accumulation in abdominal aorta tissues of CQ (40 mg/kg, i.p. LCWE-injected WT and *Sirt1^super^* mice, at 2 week post-injection (n=7,8/group). **(E)** Quantification of Western blots shown in (D). Data are presented as mean ± SEM. Each symbol represents one individual mouse. Pooled from 2 individual experiments (A, D, E) and representative of two individual experiments (B). \**p-value*<0.05, \*\**p-value*<0.01, \*\*\**p-value*<0.001, and \*\*\*\**p-value*<0.001 by unpaired t-test (B) or two-way ANOVA with Tukey post hoc test (E). CQ, chloroquine.

**Figure S7. SIRT1 overexpression inhibits LCWE-induced pathogenic phenotypic switch of VSMCs. (A)** Immunofluorescent staining and MFI of Lumican (red) in heart tissues from PBS and LCWE-injected WT and *Sirt1^super^* mice (n=5/group). DAPI (blue) is used to stain nuclei. Scale bars, 50μm. **(B)** Immunofluorescent staining and MFI of Lumican (red) in abdominal aortas of PBS- and LCWE-injected WT and *Sirt1^super^* mice at 2 weeks post-injection (n=5/group). DAPI (blue) is used to stain nuclei. Scale bars, 50μm. **(C)** Immunofluorescent staining and MFI of αSMA (green) in heart tissues from PBS or LCWE-injected WT and *Sirt1^super^*mice at 2 weeks post-injection (n=5/group). DAPI (blue) is used to stain nuclei. Scale bars, 50μm. **(D)** Immunofluorescent staining and MFI of αSMA (green) in abdominal aortas of PBS or LCWE-injected WT and *Sirt1^super^* mice at 2 weeks post-injection (n=5/group). DAPI (blue) is used to stain nuclei. Scale bars, 50μm. **(E)** mRNA levels of *Mmp9* and *Myh11* from untreated (medium), LCWE (60μg/ml, 24 hours) or recombinant (r)IL-1β (10ng/ml, 24 hours) treated primary VSMCs differentiated from abdominal aortas of WT and *Sirt1^super^* mice (n=3, 4/group). Data are presented as mean ± SEM. Each symbol represents one individual mouse. Representative of two independent experiments (A-E). \**p-value*<0.05, \*\**p-value*<0.01, \*\*\**p-value*<0.001, and \*\*\*\**p-value*<0.001 by two-way ANOVA with Tukey’s post hoc test (A, B, C, D, E). rIL-1β, recombinant IL-1β, αSMA, α-smooth muscle actin, MFI, mean fluorescence intensity.

**Figure S8. SIRT1 activation promotes autophagy/mitophagy in VSMCs *in vitro*. (A)**Quantification of band intensities of western blots (Figure 6B) relative to β-actin and LC3II/I ratio. **(B)** Quantification of band intensities of western blots (Figure 6C) relative to Ponceau S. **(C)** Quantification of band intensities of western blots (Figure 6D) relative to β-actin and LC3II/I ratio. Data are presented as mean ± SEM. Representative of two individual experiments (A-C). \**p-value*<0.05, \*\**p-value*<0.01, \*\*\**p-value*<0.001, and **** *p-value*<0.001 by one-way ANOVA with Tukey’s post hoc test (A, B, C).

**Figure S9. SIRT1 increases autophagy and reduces mitochondrial ROS formation *in vitro* (A)** Representative pictures of CytoID staining of WT primary coronary artery VSMCs s treated with SRT1720 (5μM, 24 hours) or EX527 (5μM, 24 hours) alone or in combination with rIL-1β (10ng/ml, 24 hours) and CQ (5μM, last 4 hours) (see Figure 6E) (n=3/group). Scale bars: 50 µm. **(B)** Representative mitoSOX and mitoTracker staining of WT primary coronary artery VSMCs treated with SRT1720 (5μM, 24 hours) or EX527 (5μM, 24 hours) alone or in combination with rIL-1β (10ng/ml, 24 hours) (see Figure 6F) (n=8/group). **(C)** Quantification of band intensities of western blots (Figure 6H) relative to Ponceau S. from figure 6G. Scale bars: 50 µm. Representative of 2 individual experiments. \**p-value*<0.05, \*\**p-value*<0.01, \*\*\**p-value*<0.001, and **** *p-value*<0.001 by one-way ANOVA (A-C) or two-way ANOVA with Tukey post hoc test (C).

**Figure S10. Myeloid cells specific deletion of *Sirt1* impairs autophagy and enhances M1-like macrophage polarization. (A)** Schematic of the experimental design. *Sirt1^fl/fl^* and *Myh11^Cre/ERT2^Sirt1^Δ/Δ^*mice were fed a tamoxifen diet for 2 weeks, rested for 4 weeks, and then injected with LCWE. Mice were analyzed 2 weeks after LCWE injection. **(B)** Representative western blot and quantifications of SIRT1 and β-actin in the abdominal aortas of tamoxifen-treated and LCWE-injected *Sirt1^fl/fl^*and *Myh11^Cre/ERT2^Sirt1^Δ/Δ^* mice (n=4 to 5/group). **(C)** Primary smooth muscle cells differentiated from the abdominal aortas of *Sirt1^fl/fl^* and *Myh11^Cre/ERT2^Sirt1^Δ/Δ^* mice and treated with 4OHT (1μM, 48 hours) and rIL-1β (10ng/ml, 24 hours) *in vitro*. Representative Western blots and quantifications of SIRT1, p62 LC3-II/I and β-actin levels from whole cell lysate (n=3/group). **(D)** Western blot analysis detecting SIRT1 and β-actin in BMDMs generated from *Sirt1^fl/fl^* and *Csf1r^Cre^Sirt1^Δ/Δ^*mice (n=3/group). **(E-F)** BMDMs differentiated from *Sirt1^fl/fl^*and *Csf1r^Cre^Sirt1^Δ/Δ^* mice *in vitro*. **(E)** IL-1β and TNF-α measurements in the supernatants of BMDMs from *Sirt1^fl/fl^* and *Csf1r^Cre^Sirt1^Δ/Δ^* mice unstimulated or stimulated with LCWE (60μg/ml, 24 hours) (n=6/group). **(F)** qPCR PCR measurements of mRNA levels of *Nos2, Tnfa, Fizz1 and Ym1,* normalized to *Gapdh*) in BMDMs generated from *Sirt1^fl/fl^* and *Csf1r^Cre^Sirt1^Δ/Δ^* mice after 24 hours of either LPS (10ng/ml) and IFN*γ* (50ng/ml) stimulation for M1-like polarization or IL-4 (10ng/ml) for M2 polarization (n=3,4/group). Data are presented as mean ± SEM. Each symbol represents one individual mouse in C and E. Representative of one experiment (B, C) and representative of two individual experiments (D, E, F). \**p-value*<0.05, \*\**p-value*<0.01, \*\*\**p-value*<0.001, and \*\*\*\**p-value*<0.001 by unpaired t-test (A) or two-way ANOVA with Tukey post hoc test (B, D, E). BMDM, bone marrow-derived macrophages, 4OHT, 4-hydroxytamoxifen.

**Figure S11. SIRT1 deletion in myeloid cells increases the infiltration of F4/80^+^NLRP3^+^ cells with caspase-1 activity into LCWE-induced cardiovascular lesions. (A)** Representative immunofluorescent staining of isotype, FLICA (green), F4/80 (red) and NLRP3 (purple) staining in heart tissues from PBS or LCWE injected *Sirt1^fl/fl^* and *Csf1r^Cre^Sirt1^Δ/Δ^*mice at 2 weeks post-injection (n=5 to 7/group) from Figure 7I. **(B)** Representative immunofluorescent staining of isotype, FLICA (green), F4/80 (red) and NLRP3 (purple) staining in abdominal aortas from PBS or LCWE injected *Sirt1^fl/fl^*and *Csf1r^Cre^Sirt1^Δ/Δ^* mice at 2 weeks post-injection (n=4,5 to 8/group) from Figure 7J. Pooled from 2 individual experiments (A, B). Scale bars: 50 µm. FLICA, Fluorochrome Labeled Inhibitors of Caspases.

**Figure S12. Increasing SIRT1 activity decreases the severity of cardiovascular inflammation in LCWE-induced KD. (A)** Schematic of the experimental design. WT mice were orally treated with Resveratrol (RES) by oral gavage daily, starting the day before LCWE injection until the experimental endpoint, and vascular tissues were analyzed 7 days post-LCWE injection (**B)** Representative H&E-stained heart sections and heart vessel inflammation scores of LCWE-injected WT mice untreated or orally treated with RES (n=9 to 10/group). Scale bars, 500μm. **(C, D)** Representative pictures of the abdominal aorta areas (C), maximal abdominal aorta diameter, and abdominal aorta area measurements (D) of LCWE-injected WT mice, untreated or orally treated with RES (n=9 to 10/group). **(E)** Schematic of the experimental design. WT mice were injected i.p. daily with the SIRT1 activator SRT1720, starting the day before LCWE injection until day 7. Tissues were harvested at day 14. **(F, G)** Schematic of the experimental design. WT mice received in the drinking water either β-NMN (F) or daily oral gavage of NR (G) starting the day before LCWE-injection and until day 7 post-LCWE injection, when the mice were processed. **(H)** NAD^+^ levels measured from serum of LCWE-injected WT mice that receive normal drinking water or drinking water supplemented with β-NMN at day 7 post-LCWE injection. **(I)** NAD^+^ levels measured in the serum of LCWE-injected WT mice untreated or treated daily by oral gavage with NR, at day 7 post-LCWE injection. Data are presented as mean ± SEM. Each symbol represents one individual mouse. Pooled from 2 individual experiments (B-D), representative of two experiments (H, I). \**p-value*<0.05, \*\**p-value*<0.01, \*\*\**p-value*<0.001, and \*\*\*\**p-value*<0.001 by unpaired t-test with Welch’s correction (B) and unpaired t-test (D, H, I). RES, resveratrol, NAD^+^, nicotinamide adenine dinucleotide, β-NMN, β-Nicotinamide mononucleotide, NR, nicotinamide riboside.

**Figure S13. Supplementation of NAD^+^ precursors therapeutically decreases LCWE-induced KD vasculitis. (A)** Schematic of the experimental design. WT mice were injected with LCWE, and 3 days later treated with either β-NMN in drinking water, or NR by oral gavage. Severity of LCWE-induced KD was then assessed 14 days post-LCWE injection. (**B**) Representative H&E-stained heart sections and heart vessel inflammation scores of LCWE-injected WT mice treated or not with β-NMN, at 2 weeks post-LCWE injection (n=10/group). Scale bars, 500μm. **(C, D**) Representative pictures of the abdominal aorta areas (C), maximal abdominal aorta diameter and abdominal aorta area measurements (D) of LCWE-injected WT mice treated or not with β-NMN, at 2 weeks post-LCWE injection (n=10/group). **(E)** Representative H&E-stained heart sections and heart vessel inflammation scores of LCWE-injected WT mice treated or not with NR, at 2 weeks post-LCWE injection (n=10/group). Scale bars, 500μm. **(F, G**) Representative pictures of the abdominal aorta areas (F), maximal abdominal aorta diameter and abdominal aorta area measurements (G) of LCWE-injected WT mice treated or not with NR, at 2 weeks post-LCWE injection (n=10/group). Data are presented as mean ± SEM. Each symbol represents one individual mouse. \**p-value*<0.05, \*\**p-value*<0.01, \*\*\**p-value*<0.001, and \*\*\*\**p-value*<0.001 by Mann-Whitney test (E, G) or unpaired t-test (B, D). β-NMN, β-Nicotinamide mononucleotide, NR, nicotinamide riboside.

## Notes

### Competing Interest Statement

The authors have declared no competing interest.

