## Supplementary Figures for "Sirtuin 1 Activation Mitigates Murine Vasculitis Severity by Promoting Autophagy and Mitophagy"

### Supplementary Figure 1

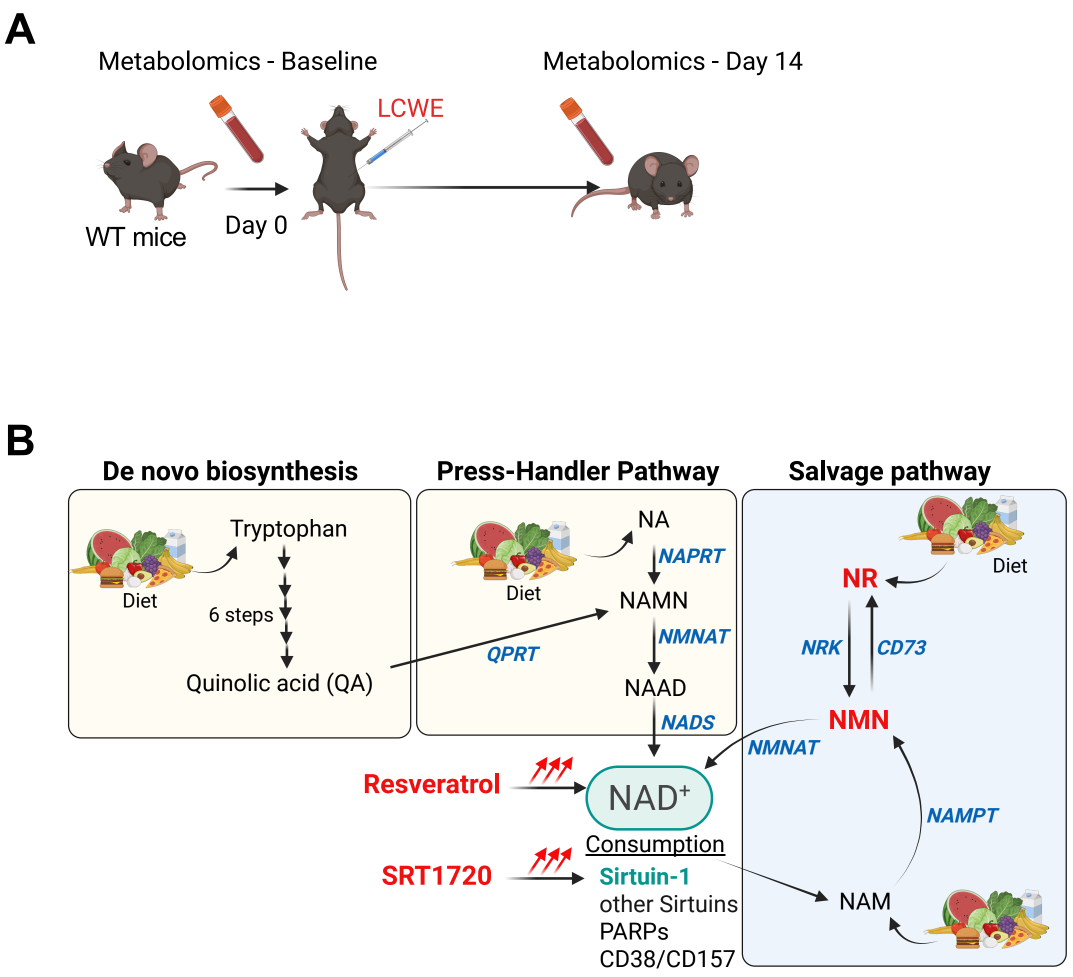


**Figure S1. Metabolomic analysis revealed a deficit in the NAD$^+$ pathway during LCWE-induced KD. (A)** Schematic of the experimental design. Blood was collected from WT mice at day 0 (baseline) and 14 days after LCWE injection for metabolomics analysis. (**B**) Schematic of the different pathways involved in NAD$^+$ production, depletion, and consumption. NAD$^+$, nicotinamide adenine dinucleotide, NR, nicotinamide riboside, NMN, nicotinamide mononucleotide, NAM, nicotinamide, NRK, nicotinamide riboside kinase, NAMPT, nicotinamide phosphoribosyltransferase, NAMNAT, nicotinamide/nicotinate mononucleotide adenylyltransferase, QA, quinolinic acid, QPRT, quinolinate phosphoribosyl transferase, NA, nicotinic acid, NAMN, nicotinic acid mononucleotide, NAPRT, NA phosphoribosyltransferase, NMNAT, NMN adenylyltransferases, NADS, NAD synthase, PARPs, poly (ADP-ribose) polymerases.

#### Supplementary Figure 2

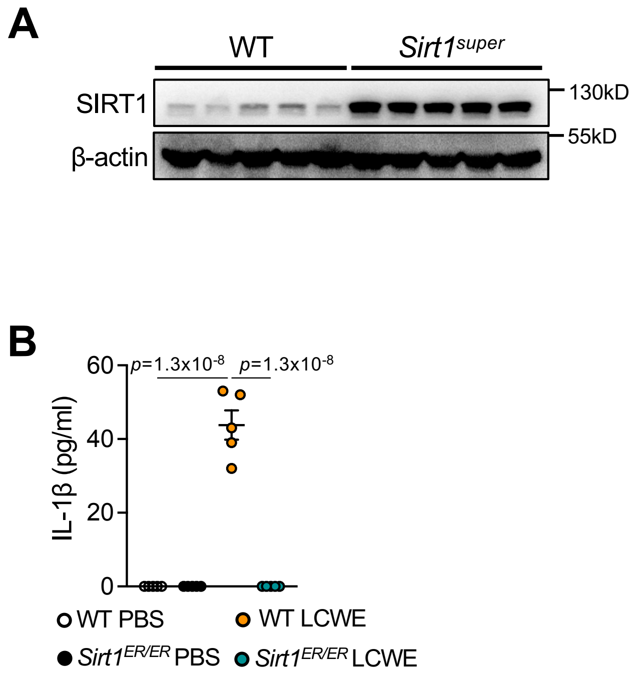


**Figure S2. SIRT1 protein levels in heart tissues of transgenic mice. (A)** Protein lysates of the apex part of heart tissues from WT and *Sirt1*$^{super}$ mice injected with LCWE were analyzed by Western blotting using specific antibodies for SIRT1 and β-actin (n=5/group). Representative of 2 experiments. **(B)** Levels of IL-1β in the peritoneal lavage of WT and *Sirt1*$^{ER/ER}$ mice injected 5 consecutive days of tamoxifen (75mg/kg) and then with PBS or LCWE 24 hours post-injection (n=5/group). **p-value*<0.05, ***p-value*<0.01, ****p-value*<0.001, and *****p-value*<0.001 by two-way ANOVA with Tukey post hoc test (B).

### Supplementary Figure 3

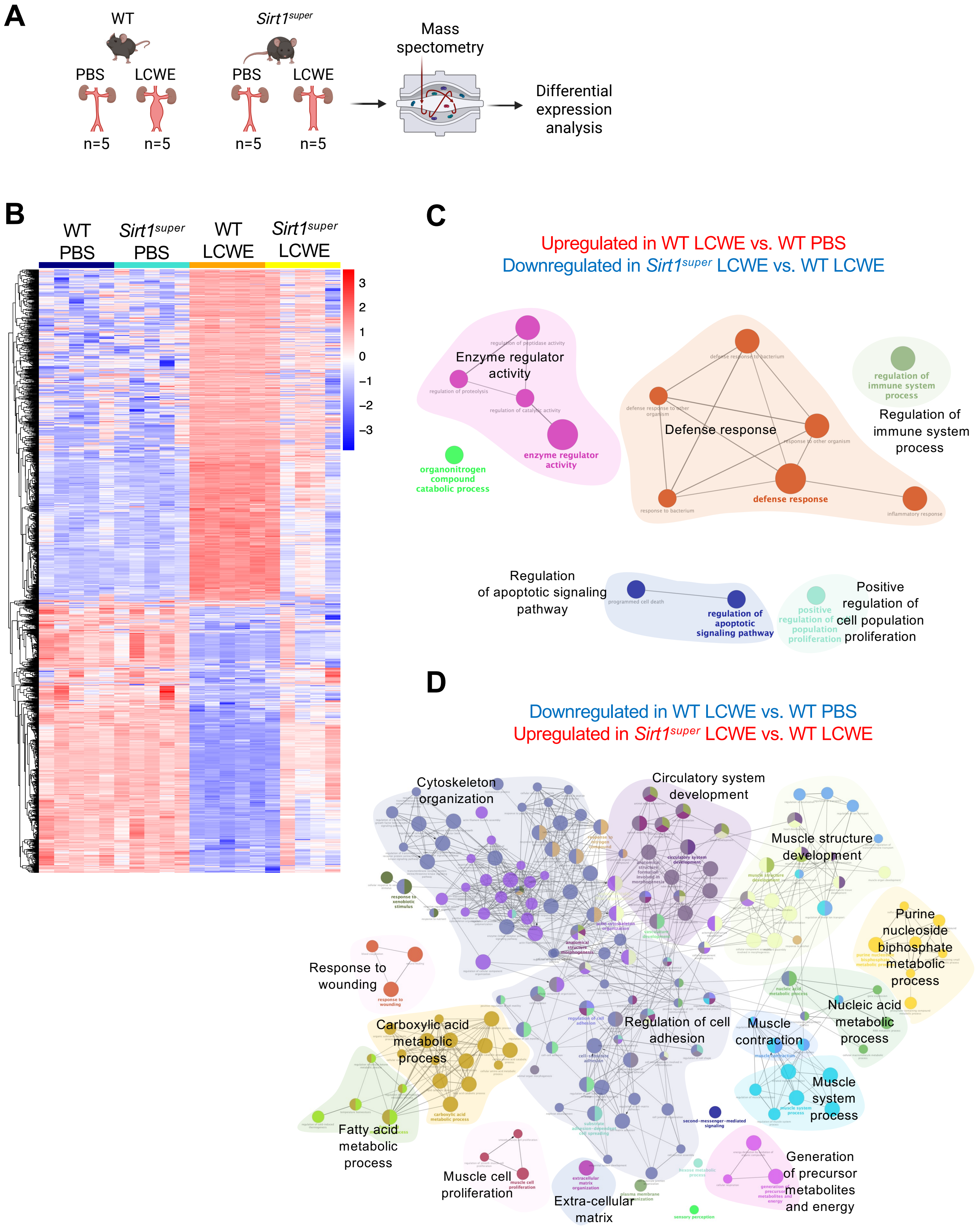

**Figure S3. SIRT1 overexpression counteracts LCWE-induced KD related inflammation in abdominal aortas. (A)** Schematic of the experimental design. Abdominal aortas of either PBS or LCWE-injected WT and *Sirt1*$^{super}$ mice were collected 2 weeks post-injection (n=5/group), and tissue processed for untargeted proteomics analysis. **(B)** Heatmap of global protein expression data from abdominal aortas of either PBS or LCWE-injected WT and *Sirt1*$^{super}$ mice 2 weeks post-injection (n=5/group). **(C)** ClueGO Ontology analysis of pathways from LCWE-induced KD DEPs the highlighted proteins in red from (B), which are significantly increased in LCWE-injected WT mice compared with PBS-injected WT mice and downregulated in LCWE-injected *Sirt1*$^{super}$ mice compared with LCWE-injected WT mice (n=5/group). **(D)** ClueGO Ontology analysis of pathways from the highlighted proteins in red from (B), which are significantly decreased in LCWE-injected WT mice compared with PBS-injected WT mice and increased in LCWE-injected *Sirt1*$^{super}$ mice compared with LCWE-injected WT mice (n=5/group).

### Supplementary Figure 4

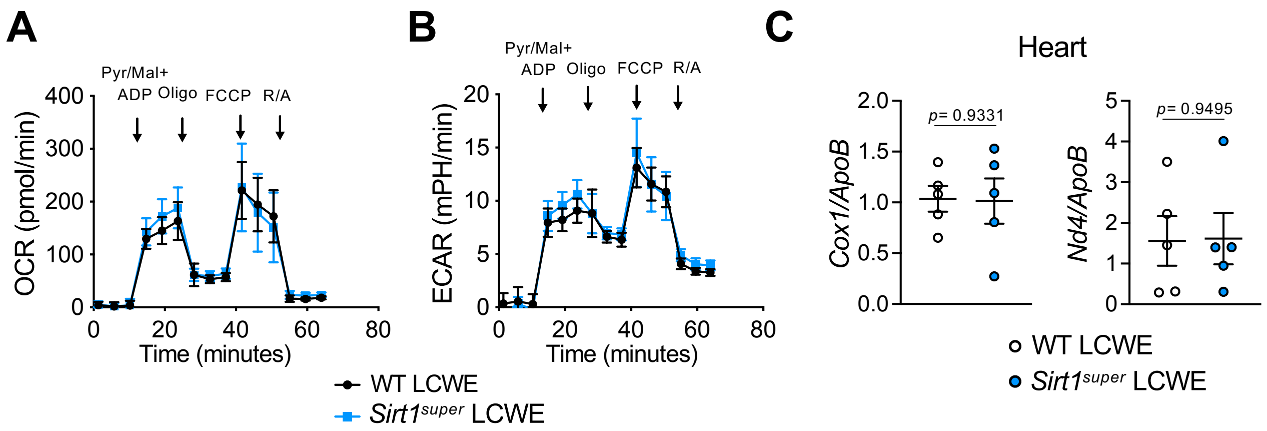


**Figure S4. SIRT1 increases mitochondrial respiration efficiency but does not impact mtDNA levels during LCWE-induced KD (A)** Oxygen consumption rate (OCR) in isolated mitochondria of heart tissues from LCWE-injected WT or *Sirt1*$^{super}$ mice at 2 weeks post-injection. (n=5/group). **(B)** Extracellular acidification rate (ECAR) in isolated mitochondria of heart tissues from LCWE-injected WT or *Sirt1*$^{super}$ mice at 2 weeks post-injection. (n=5/group). Oligomycin: ATP synthase inhibitor; FCCP: mitochondrial uncoupler; R/A: rotenone and antimycin A mix (inhibitors for ETC complex **I** and **III**, respectively). **(C)** qRT-PCR quantification of total genomic DNA and mtDNA (*Cox1* and *Nd4*) to nucDNA (*ApoB*) ratio from heart tissues of WT and *Sirt1*$^{super}$ mice injected with LCWE 2 weeks post-injection (n=5/group). Data are presented as mean ± SEM. Each symbol represents one individual mouse. **p-value*<0.05, ***p-value*<0.01, ****p-value*<0.001, and *****p-value*<0.001 by unpaired t-test (C). Pyr/Mal, pyruvate/malate, ADP, adenosine diphosphate, FCCP, Carbonyl cyanide p-trifluoromethoxyphenylhydrazone.

#### Supplementary Figure 5

**A** **Autophagy/ROS Metabolism**

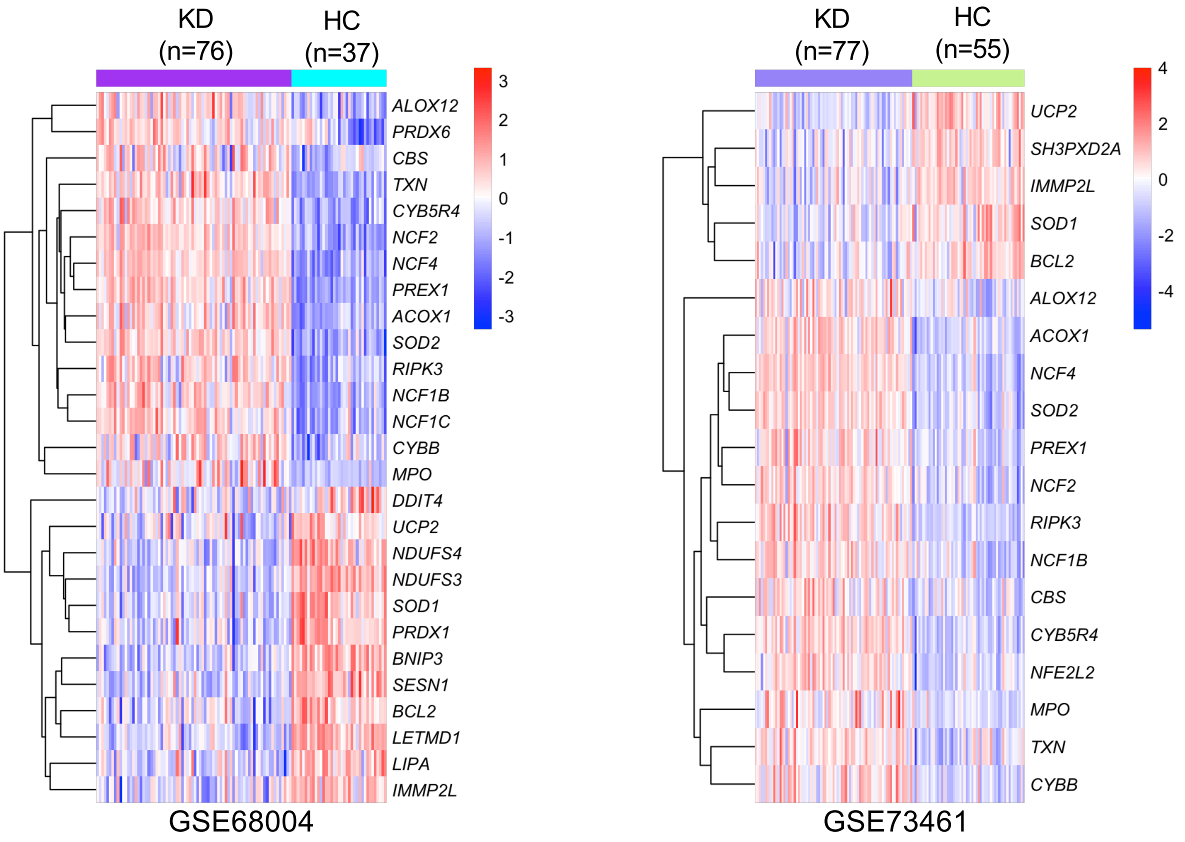


**B** **Oxidative Phosphorylation**

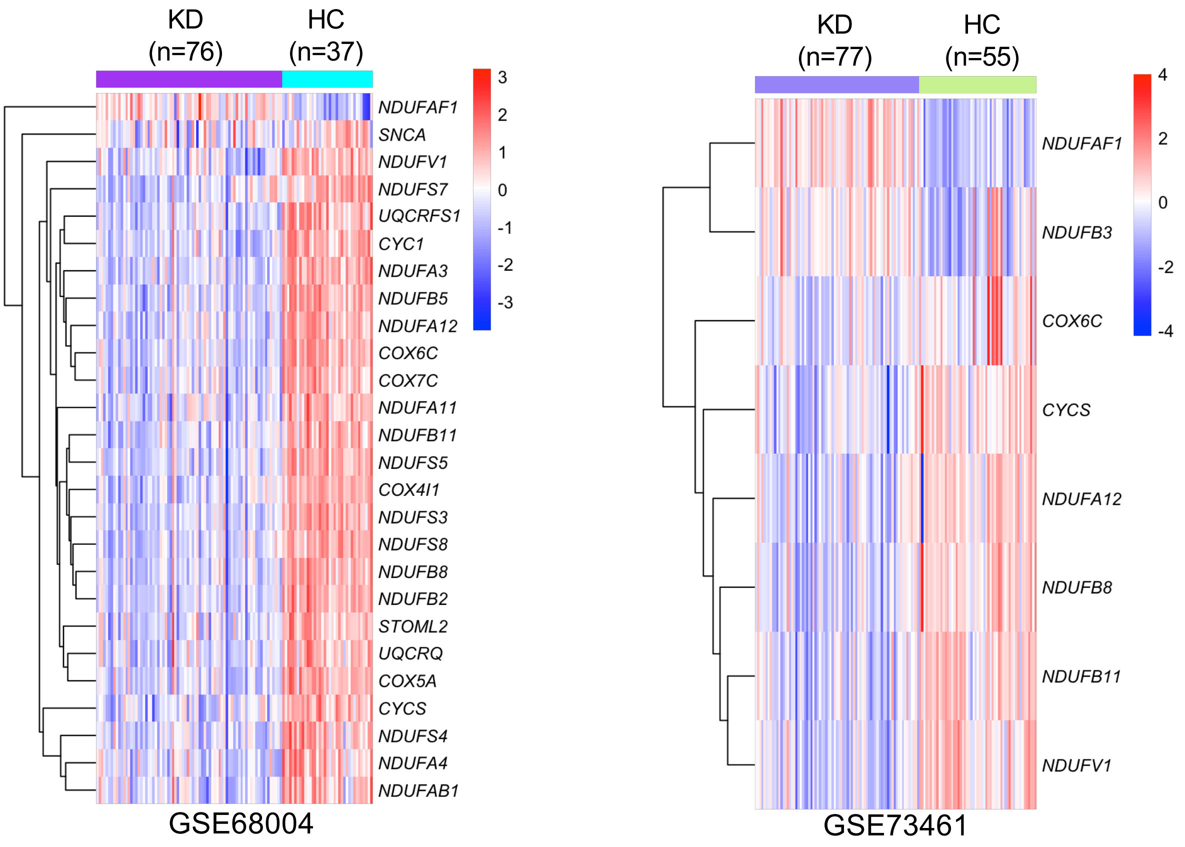

**Figure S5. Expression of signature genes involved ROS metabolism and oxidative phosphorylation in patients with KD. (A-B)** Heatmaps illustrating the expression of genes involved in ROS metabolism (A) and oxidative phosphorylation (B) in whole blood from patients with KD compared with healthy control (HC) (GSE68004 and GSE73461).

#### Supplementary Figure 6

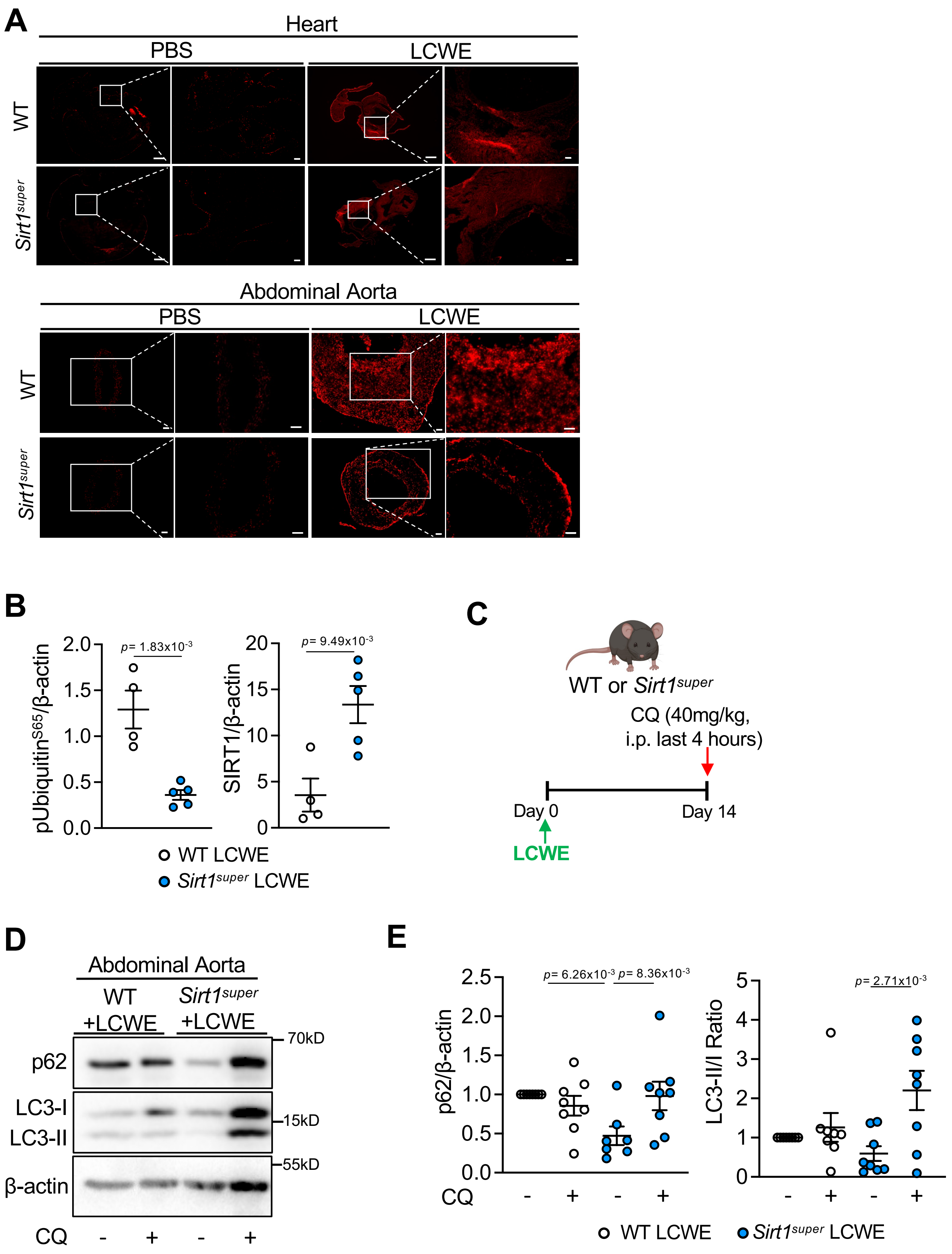

**Figure S6. Overexpression of SIRT1 reduces ROS accumulation and promotes autophagic flux during LCWE-induced KD. (A)** Representative dihydroethidium (DHE) staining heart (upper) and abdominal aorta (lower) tissue cross-sections collected from WT and *Sirt1*$^{super}$ mice injected with PBS or LCWE at 2 weeks post-injection (n=5/group). Scale bars: 50 μm. **(B)** Western blot quantification of SIRT1 and pUbiquitin$^{S65}$ in protein lysates of heart tissues from WT and *Sirt1*$^{super}$ mice injected with LCWE at 2 weeks post-injection (n=4, 5/group). **(C)** Schematic of the *in vivo* autophagic flux assay experimental design. WT and *Sirt1*$^{super}$ mice were injected with LCWE, and 14 days later received chloroquine (CQ) i.p. 4 hours before collecting the abdominal aortas. **(D)** Representative Western blots of p62, LC3-I and LC3-II protein accumulation in abdominal aorta tissues of CQ (40 mg/kg, i.p. LCWE-injected WT and *Sirt1*$^{super}$ mice, at 2 week post-injection (n=7,8/group). **(E)** Quantification of Western blots shown in (D). Data are presented as mean ± SEM. Each symbol represents one individual mouse. Pooled from 2 individual experiments (A, D, E) and representative of two individual experiments (B). **p-value*<0.05, ***p-value*<0.01, ****p-value*<0.001, and *****p-value*<0.001 by unpaired t-test (B) or two-way ANOVA with Tukey post hoc test (E). CQ, chloroquine.

### Supplementary Figure 7

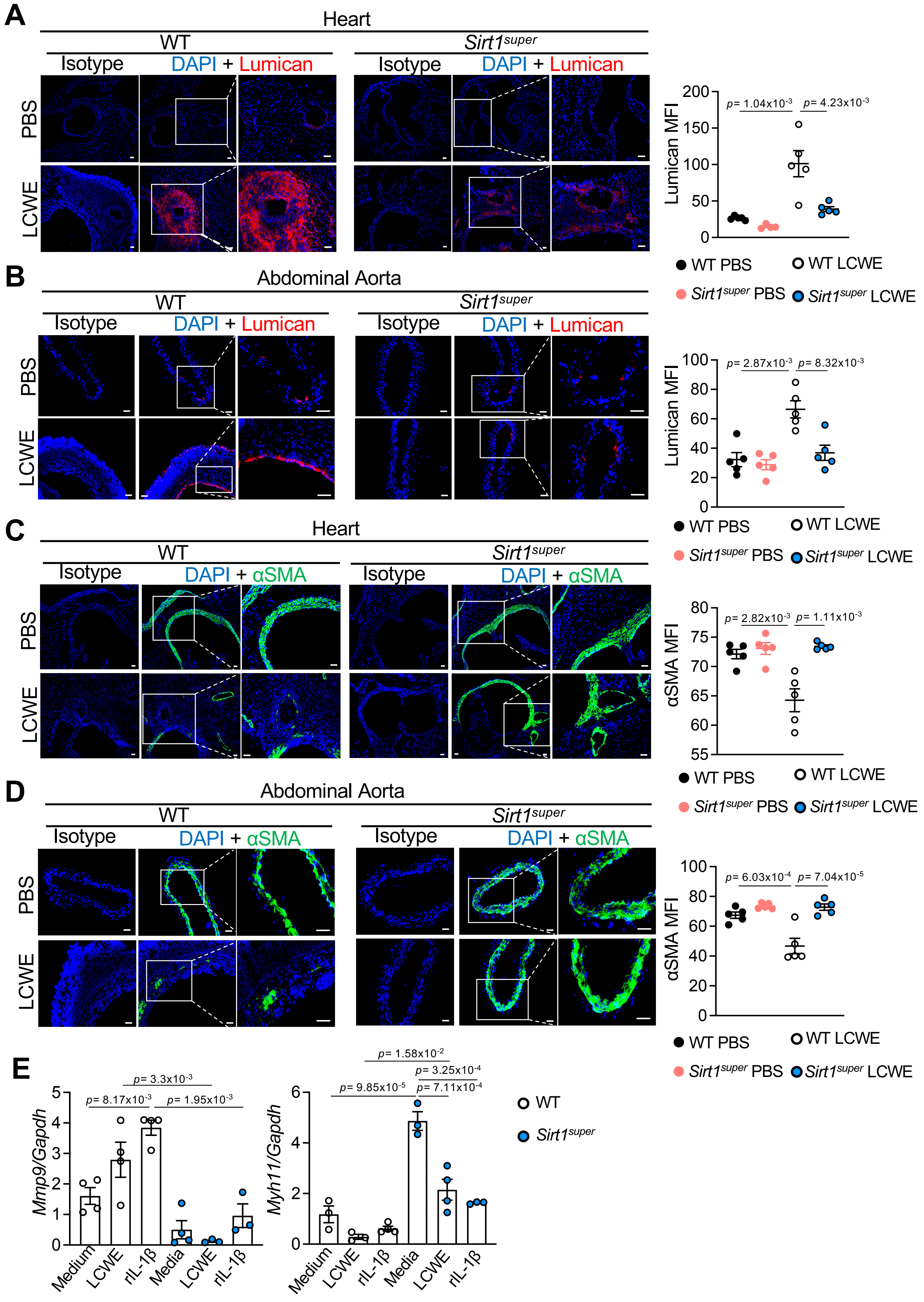

**Figure S7. SIRT1 overexpression inhibits LCWE-induced pathogenic phenotypic switch of VSMCs. (A)** Immunofluorescent staining and MFI of Lumican (red) in heart tissues from PBS and LCWE-injected WT and *Sirt1*$^{super}$ mice (n=5/group). DAPI (blue) is used to stain nuclei. Scale bars, 50µm. **(B)** Immunofluorescent staining and MFI of Lumican (red) in abdominal aortas of PBS- and LCWE-injected WT and *Sirt1*$^{super}$ mice at 2 weeks post-injection (n=5/group). DAPI (blue) is used to stain nuclei. Scale bars, 50µm. **(C)** Immunofluorescent staining and MFI of αSMA (green) in heart tissues from PBS or LCWE-injected WT and *Sirt1*$^{super}$ mice at 2 weeks post-injection (n=5/group). DAPI (blue) is used to stain nuclei. Scale bars, 50µm. **(D)** Immunofluorescent staining and MFI of αSMA (green) in abdominal aortas of PBS or LCWE-injected WT and *Sirt1*$^{super}$ mice at 2 weeks post-injection (n=5/group). DAPI (blue) is used to stain nuclei. Scale bars, 50µm. **(E)** mRNA levels of *Mmp9* and *Myh11* from untreated (medium), LCWE (60µg/ml, 24 hours) or recombinant (r)IL-1β (10ng/ml, 24 hours) treated primary VSMCs differentiated from abdominal aortas of WT and *Sirt1*$^{super}$ mice (n=3, 4/group). Data are presented as mean ± SEM. Each symbol represents one individual mouse. Representative of two independent experiments (A-E). **p-value*<0.05, ***p-value*<0.01, ****p-value*<0.001, and *****p-value*<0.001 by two-way ANOVA with Tukey's post hoc test (A, B, C, D, E). rIL-1β, recombinant IL-1β, αSMA, α-smooth muscle actin, MFI, mean fluorescence intensity.

### Supplementary Figure 8

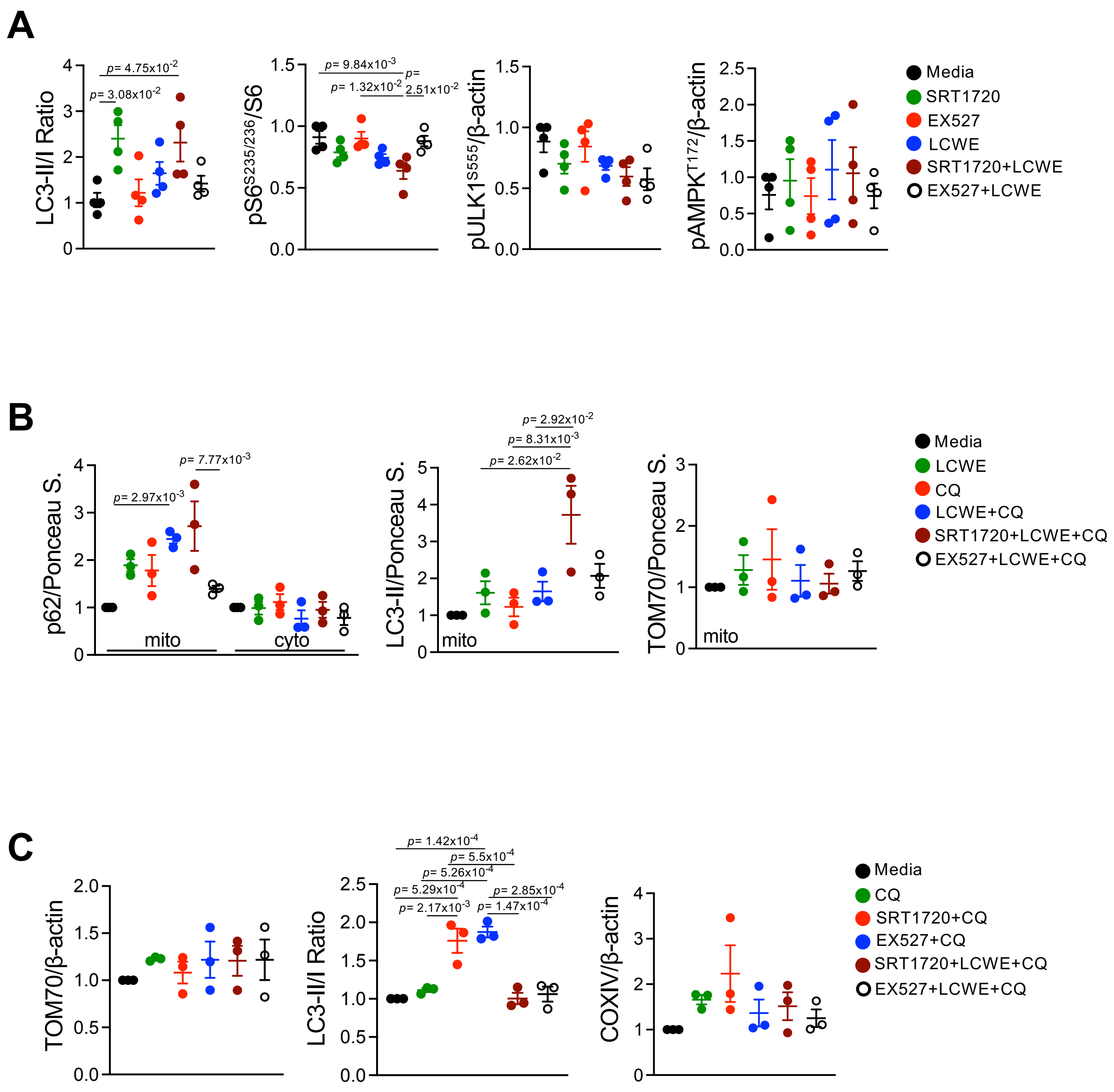


**Figure S8. SIRT1 activation promotes autophagy/mitophagy in VSMCs *in vitro*. (A)** Quantification of band intensities of western blots (Figure 6B) relative to β-actin and LC3II/I ratio. **(B)** Quantification of band intensities of western blots (Figure 6C) relative to Ponceau S. **(C)** Quantification of band intensities of western blots (Figure 6D) relative to β-actin and LC3II/I ratio. Data are presented as mean ± SEM. Representative of two individual experiments (A-C). **p-value*<0.05, ***p-value*<0.01, ****p-value*<0.001, and **** *p-value*<0.001 by one-way ANOVA with Tukey's post hoc test (A, B, C).

#### Supplementary Figure 9

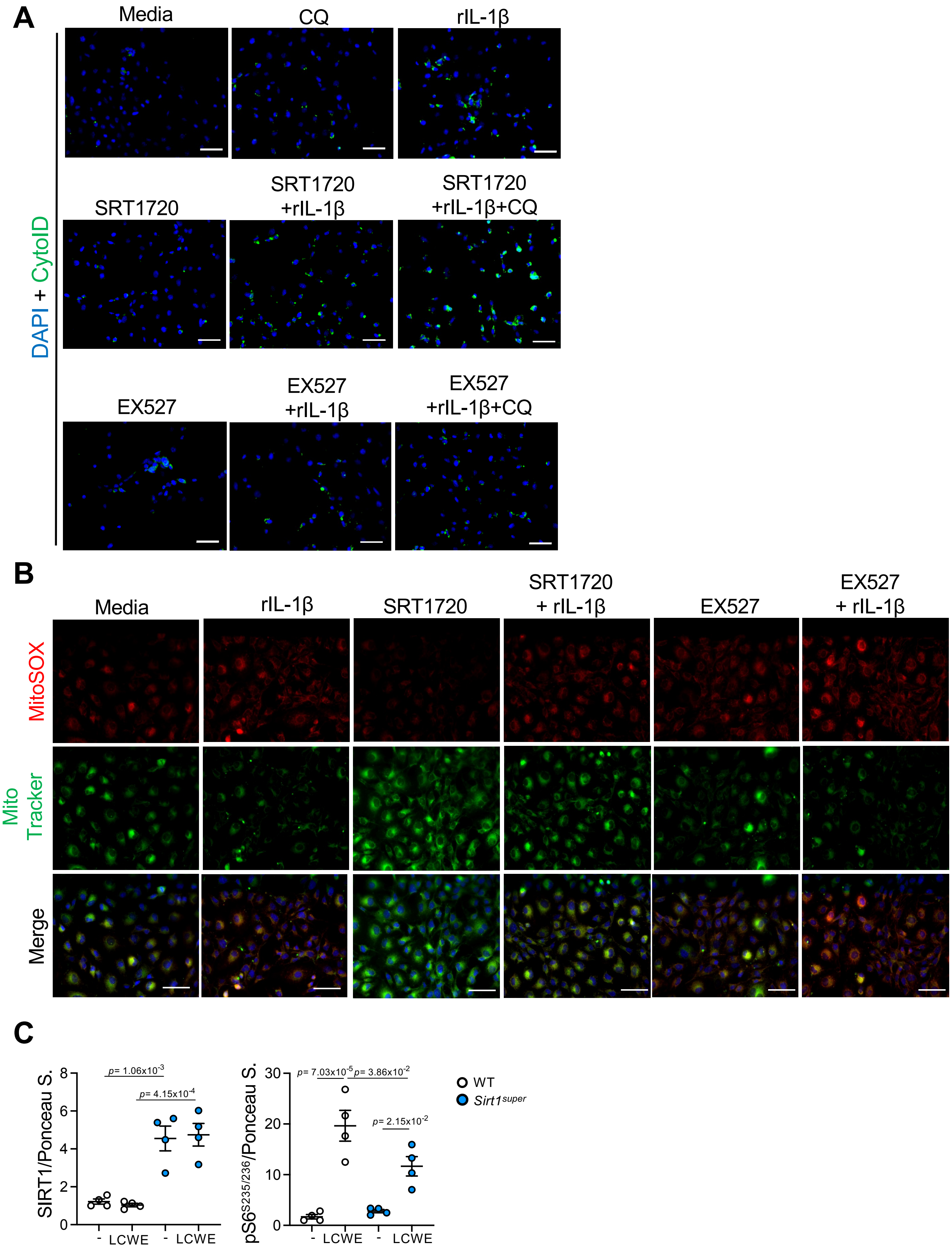

**Figure S9. SIRT1 increases autophagy and reduces mitochondrial ROS formation *in vitro* (A)** Representative pictures of CytoID staining of WT primary coronary artery VSMCs s treated with SRT1720 (5µM, 24 hours) or EX527 (5µM, 24 hours) alone or in combination with rIL-1β (10ng/ml, 24 hours) and CQ (5µM, last 4 hours) (see Figure 6E) (n=3/group). Scale bars: 50 µm. **(B)** Representative mitoSOX and mitoTracker staining of WT primary coronary artery VSMCs treated with SRT1720 (5µM, 24 hours) or EX527 (5µM, 24 hours) alone or in combination with rIL-1β (10ng/ml, 24 hours) (see Figure 6F) (n=8/group). **(C)** Quantification of band intensities of western blots (Figure 6H) relative to Ponceau S. from figure 6G. Scale bars: 50 µm. Representative of 2 individual experiments. **p-value*<0.05, ***p-value*<0.01, ****p-value*<0.001, and **** *p-value*<0.001 by one-way ANOVA (A-C) or two-way ANOVA with Tukey post hoc test (C).

### Supplementary Figure 10

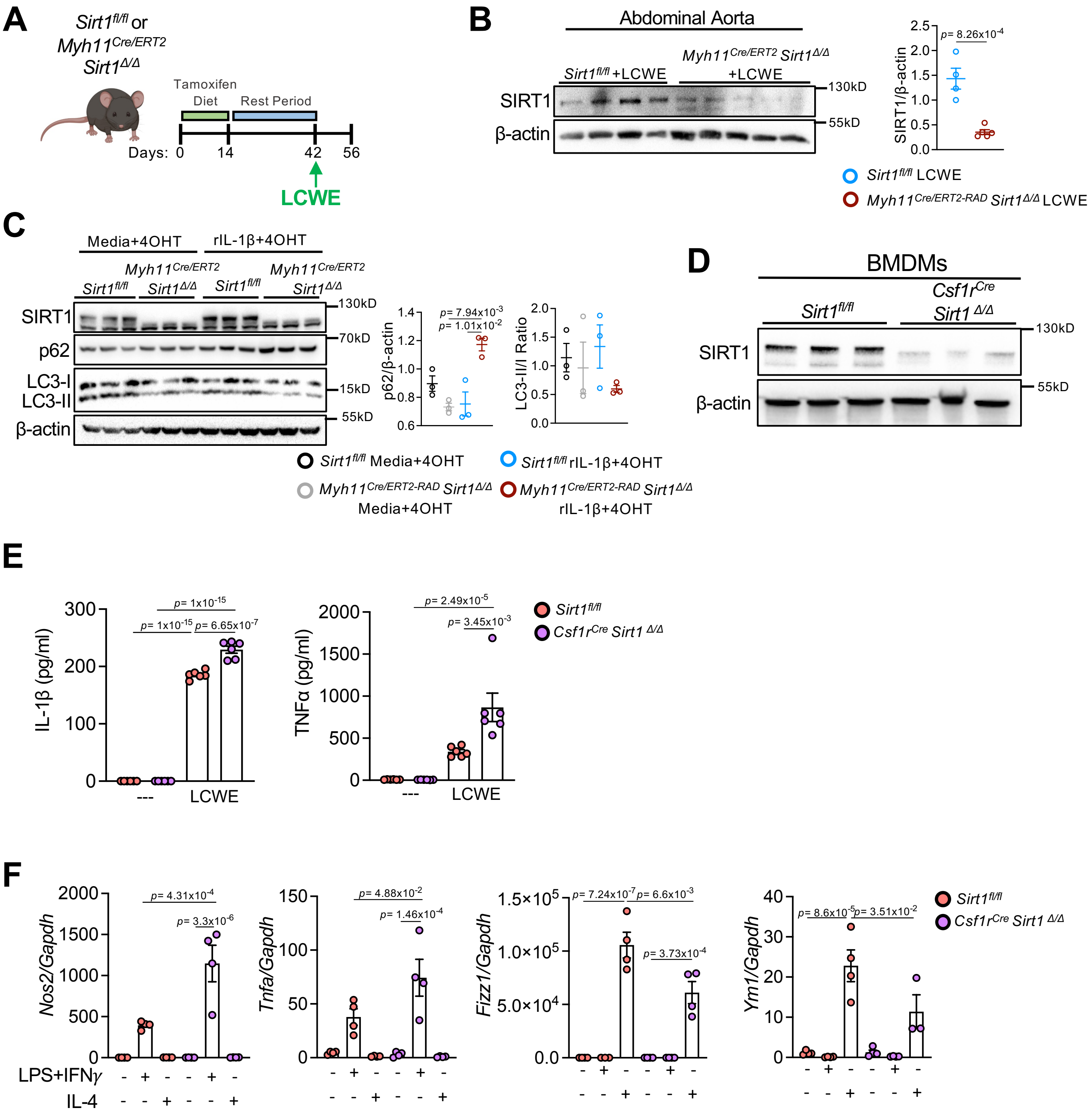

**Figure S10. Myeloid cells specific deletion of *Sirt1* impairs autophagy and enhances M1-like macrophage polarization. (A)** Schematic of the experimental design. *Sirt1*$^{fl/fl}$ and *Myh11*$^{Cre/ERT2}$*Sirt1*$^{\Delta/\Delta}$ mice were fed a tamoxifen diet for 2 weeks, rested for 4 weeks, and then injected with LCWE. Mice were analyzed 2 weeks after LCWE injection. **(B)** Representative western blot and quantifications of SIRT1 and β-actin in the abdominal aortas of tamoxifen-treated and LCWE-injected *Sirt1*$^{fl/fl}$ and *Myh11*$^{Cre/ERT2}$*Sirt1*$^{\Delta/\Delta}$ mice (n=4 to 5/group). **(C)** Primary smooth muscle cells differentiated from the abdominal aortas of *Sirt1*$^{fl/fl}$ and *Myh11*$^{Cre/ERT2}$*Sirt1*$^{\Delta/\Delta}$ mice and treated with 4OHT (1μM, 48 hours) and rIL-1β (10ng/ml, 24 hours) *in vitro*. Representative Western blots and quantifications of SIRT1, p62 LC3-II/I and β-actin levels from whole cell lysate (n=3/group). **(D)** Western blot analysis detecting SIRT1 and β-actin in BMDMs generated from *Sirt1*$^{fl/fl}$ and *Csf1r*$^{Cre}$*Sirt1*$^{\Delta/\Delta}$ mice (n=3/group). **(E-F)** BMDMs differentiated from *Sirt1*$^{fl/fl}$ and *Csf1r*$^{Cre}$*Sirt1*$^{\Delta/\Delta}$ mice *in vitro*. **(E)** IL-1β and TNF-α measurements in the supernatants of BMDMs from *Sirt1*$^{fl/fl}$ and *Csf1r*$^{Cre}$*Sirt1*$^{\Delta/\Delta}$ mice unstimulated or stimulated with LCWE (60μg/ml, 24 hours) (n=6/group). **(F)** qPCR PCR measurements of mRNA levels of *Nos2, Tnfa, Fizz1 and Ym1,* normalized to *Gapdh*) in BMDMs generated from *Sirt1*$^{fl/fl}$ and *Csf1r*$^{Cre}$*Sirt1*$^{\Delta/\Delta}$ mice after 24 hours of either LPS (10ng/ml) and IFN$\gamma$ (50ng/ml) stimulation for M1-like polarization or IL-4 (10ng/ml) for M2 polarization (n=3,4/group). Data are presented as mean ± SEM. Each symbol represents one individual mouse in C and E. Representative of one experiment (B, C) and representative of two individual experiments (D, E, F). **p-value*<0.05, ***p-value*<0.01, ****p-value*<0.001, and *****p-value*<0.001 by unpaired t-test (A) or two-way ANOVA with Tukey post hoc test (B, D, E). BMDM, bone marrow-derived macrophages, 4OHT, 4-hydroxytamoxifen.

### Supplementary Figure 11

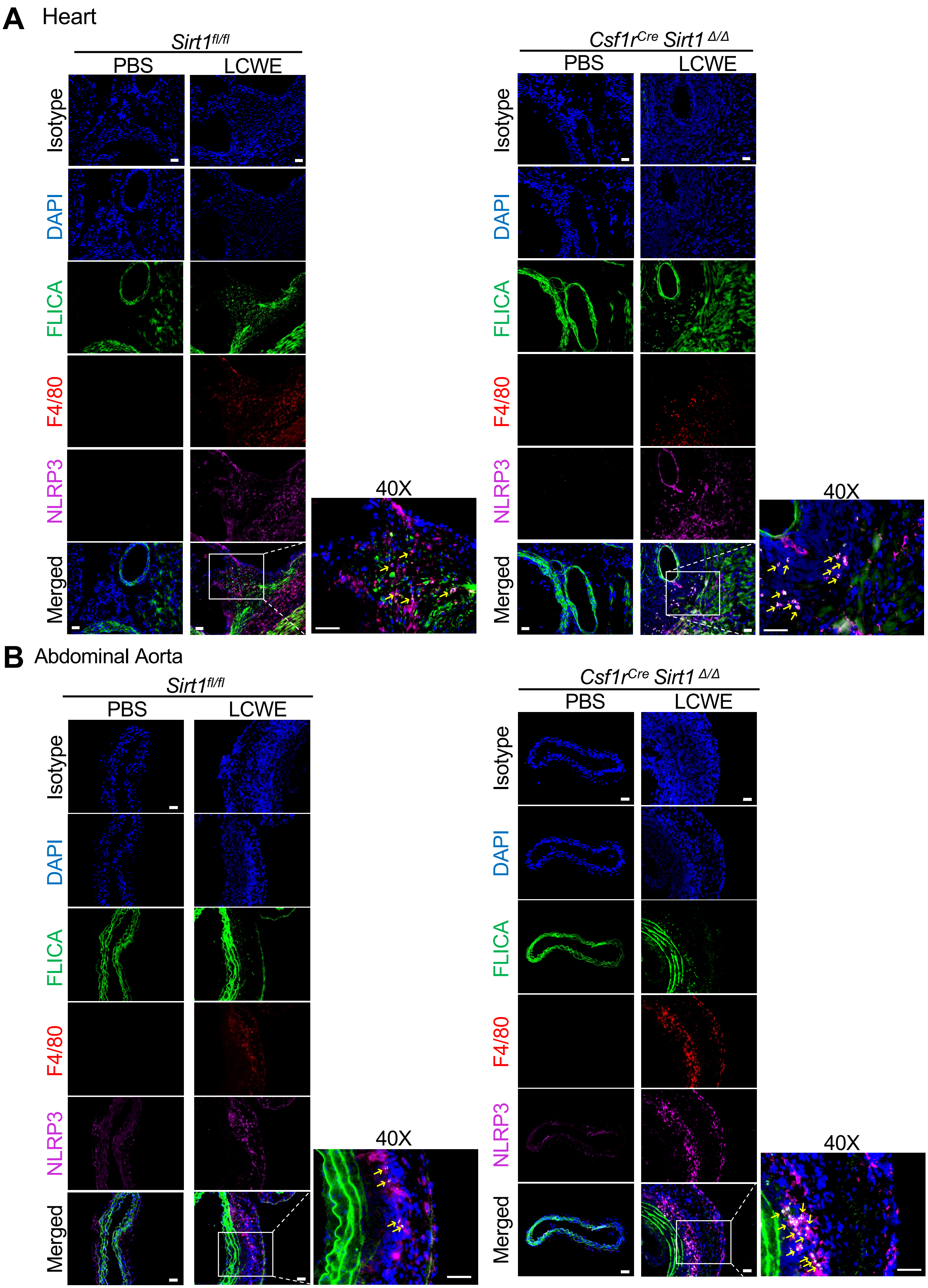

**Figure S11. SIRT1 deletion in myeloid cells increases the infiltration of F4/80$^+$ NLRP3$^+$ cells with caspase-1 activity into LCWE-induced cardiovascular lesions. (A)** Representative immunofluorescent staining of isotype, FLICA (green), F4/80 (red) and NLRP3 (purple) staining in heart tissues from PBS or LCWE injected *Sirt1$^{fl/fl}$* and *Csf1r$^{Cre}$Sirt1$^{\Delta/\Delta}$* mice at 2 weeks post-injection (n=5 to 7/group) from Figure 7I. **(B)** Representative immunofluorescent staining of isotype, FLICA (green), F4/80 (red) and NLRP3 (purple) staining in abdominal aortas from PBS or LCWE injected *Sirt1$^{fl/fl}$* and *Csf1r$^{Cre}$Sirt1$^{\Delta/\Delta}$* mice at 2 weeks post-injection (n=4,5 to 8/group) from Figure 7J. Pooled from 2 individual experiments (A, B). Scale bars: 50 µm. FLICA, Fluorochrome Labeled Inhibitors of Caspases.

### Supplementary Figure 12

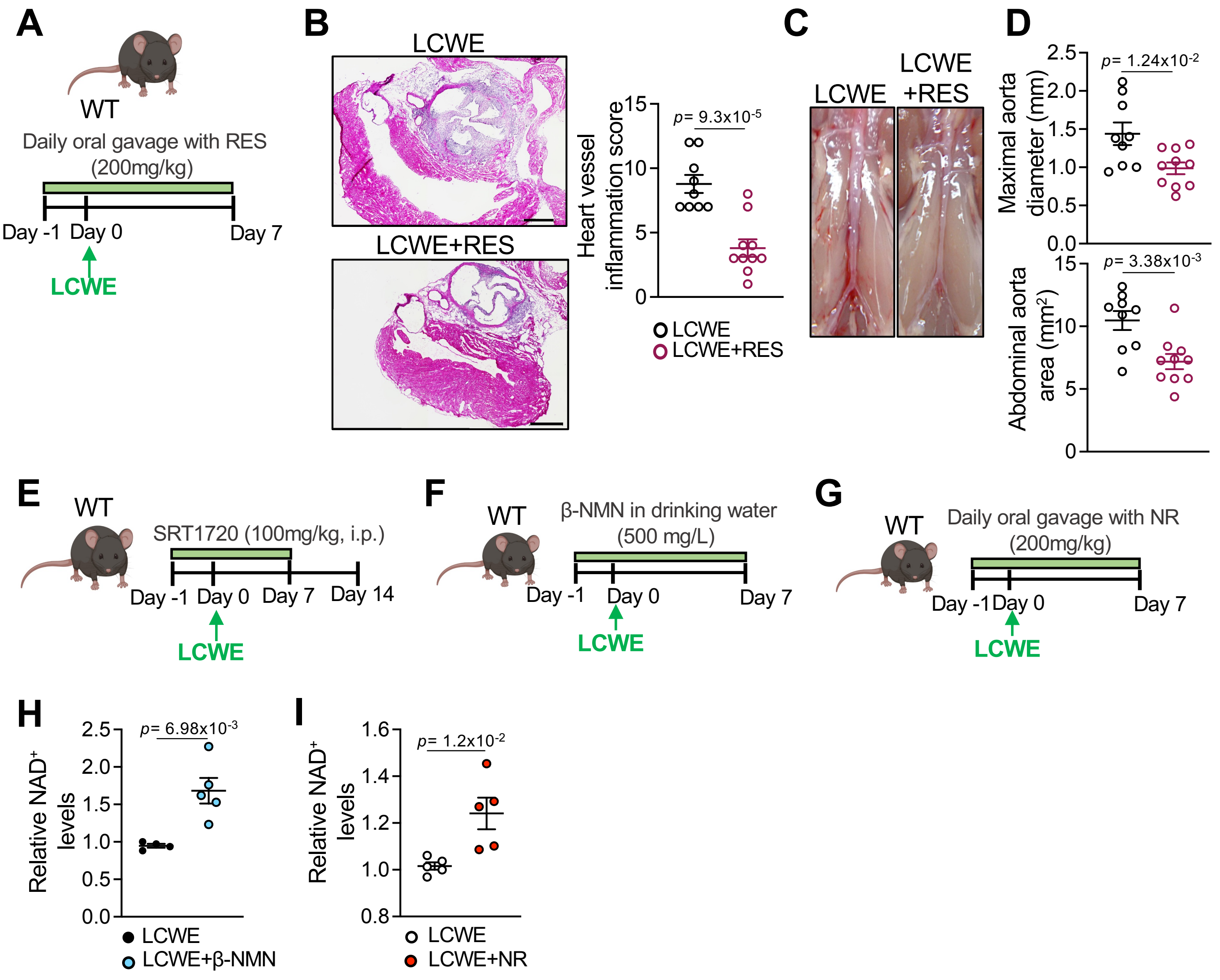


**Figure S12. Increasing SIRT1 activity decreases the severity of cardiovascular inflammation in LCWE-induced KD. (A)** Schematic of the experimental design. WT mice were orally treated with Resveratrol (RES) by oral gavage daily, starting the day before LCWE injection until the experimental endpoint, and vascular tissues were analyzed 7 days post-LCWE injection (**B**) Representative H&E-stained heart sections and heart vessel inflammation scores of LCWE-injected WT mice untreated or orally treated with RES (n=9 to 10/group). Scale bars, 500µm. **(C, D)** Representative pictures of the abdominal aorta areas (C), maximal abdominal aorta diameter, and abdominal aorta area measurements (D) of LCWE-injected WT mice, untreated or orally treated with RES (n=9 to 10/group). **(E)** Schematic of the experimental design. WT mice were injected i.p. daily with the SIRT1 activator SRT1720, starting the day before LCWE injection until day 7. Tissues were harvested at day 14. **(F, G)** Schematic of the experimental design. WT mice received in the drinking water either β-NMN (F) or daily oral gavage of NR (G) starting the day before LCWE-injection and until day 7 post-LCWE injection, when the mice were processed. **(H)** NAD$^+$ levels measured from serum of LCWE-injected WT mice that receive normal drinking water or drinking water supplemented with β-NMN at day 7 post-LCWE injection. **(I)** NAD$^+$ levels measured in the serum of LCWE-injected WT mice untreated or treated daily by oral gavage with NR, at day 7 post-LCWE injection. Data are presented as mean ± SEM. Each symbol represents one individual mouse. Pooled from 2 individual experiments (B-D), representative of two experiments (H, I). **p-value*<0.05, ***p-value*<0.01, ****p-value*<0.001, and *****p-value*<0.001 by unpaired t-test with Welch's correction (B) and unpaired t-test (D, H, I). RES, resveratrol, NAD$^+$, nicotinamide adenine dinucleotide, β-NMN, β-Nicotinamide mononucleotide, NR, nicotinamide riboside.

### Supplementary Figure 13

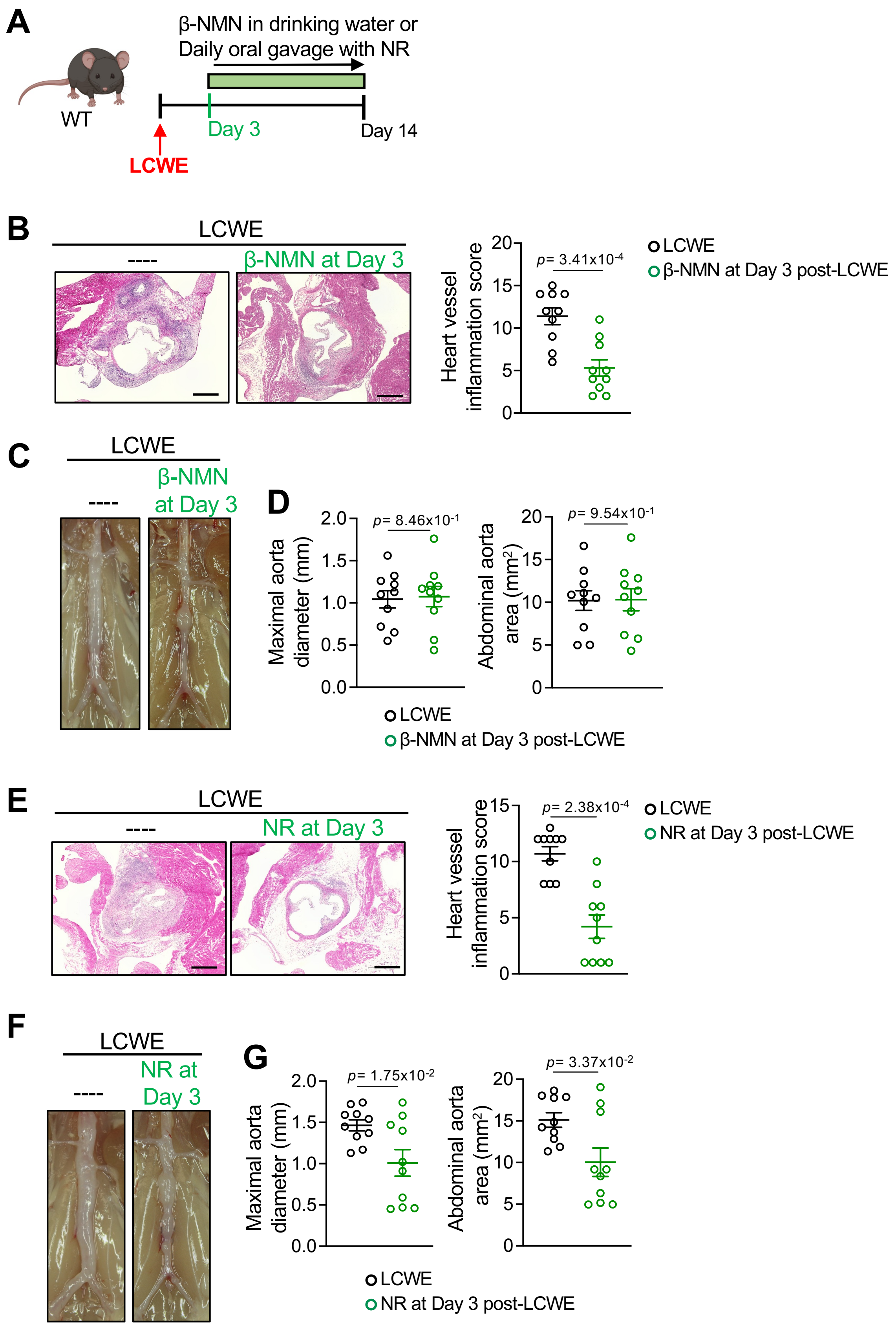

**Figure S13. Supplementation of $NAD^+$ precursors therapeutically decreases LCWE-induced KD vasculitis. (A)** Schematic of the experimental design. WT mice were injected with LCWE, and 3 days later treated with either β-NMN in drinking water, or NR by oral gavage. Severity of LCWE-induced KD was then assessed 14 days post-LCWE injection. (**B**) Representative H&E-stained heart sections and heart vessel inflammation scores of LCWE-injected WT mice treated or not with β-NMN, at 2 weeks post-LCWE injection (n=10/group). Scale bars, 500µm. **(C, D**) Representative pictures of the abdominal aorta areas (C), maximal abdominal aorta diameter and abdominal aorta area measurements (D) of LCWE-injected WT mice treated or not with β-NMN, at 2 weeks post-LCWE injection (n=10/group). **(E)** Representative H&E-stained heart sections and heart vessel inflammation scores of LCWE-injected WT mice treated or not with NR, at 2 weeks post-LCWE injection (n=10/group). Scale bars, 500µm. **(F, G**) Representative pictures of the abdominal aorta areas (F), maximal abdominal aorta diameter and abdominal aorta area measurements (G) of LCWE-injected WT mice treated or not with NR, at 2 weeks post-LCWE injection (n=10/group). Data are presented as mean ± SEM. Each symbol represents one individual mouse. **p-value*<0.05, ***p-value*<0.01, ****p-value*<0.001, and *****p-value*<0.001 by Mann-Whitney test (E, G) or unpaired t-test (B, D). β-NMN, β-Nicotinamide mononucleotide, NR, nicotinamide riboside.
